# Loss of C9orf72 perturbs the Ran-GTPase gradient and generates compositionally diverse cytoplasmic Importin β-1 granules in motor and cortical neurons *in vivo*

**DOI:** 10.1101/2021.10.20.465148

**Authors:** Philip McGoldrick, Agnes Lau, Zhipeng You, Thomas M Durcan, Janice Robertson

**Author notes:** Correspondence should be addressed to: Janice Robertson, Tanz Centre for Neurodegenerative Diseases, 60 Leonard Avenue, Toronto, ON, Canada, M5T 2S8, Philip McGoldrick, Tanz Centre for Neurodegenerative Diseases, 60 Leonard Avenue, Toronto, ON, Canada, M5T 2S8.

## Abstract

Repeat expansions in *C9orf72* cause Amyotrophic Lateral Sclerosis (ALS) and Frontotemporal Dementia (FTD) eliciting toxic effects through generation of RNA foci, dipeptide repeat proteins and/or loss of C9orf72 protein. Defects in nucleocytoplasmic transport (NCT) have been implicated as a pathogenic mechanism underlying repeat expansion toxicity. Here, we show that loss of C9orf72 causes neuronal specific phenotypes, disrupting the Ran-GTPase gradient both *in vitro* and *in vivo*. We describe compositionally different types of cytoplasmic Importin β-1 granules that exhibit neuronal subtype-specific properties *in vivo*. We show that the abundance of Importin β-1 granules is increased in the context of C9orf72 deficiency, disrupting interactions with nuclear pore complex proteins. These granules appear to bud from the nuclear envelope and are co-immunoreactive for G3BP1 and K63-ubiquitin. These findings link loss of C9orf72 protein to gain-of-function mechanisms and defects in NCT.

## Introduction

A GGGGCC (G_4_C_2_) hexanucleotide repeat expansion in the first intron of *C9orf72* (chromosome 9 open reading frame 72) is the most frequent genetic cause of amyotrophic lateral sclerosis (ALS) and frontotemporal dementia (FTD)^1,2^, two adult-onset neurodegenerative diseases entwined by overlapping clinical, genetic and neuropathological features. Three non-mutually exclusive mechanisms have been associated with the *C9orf72* mutation: RNA toxicity caused by sequestration of RNA-binding proteins to repeat expansion-containing RNA foci^1^, deleterious effects of dipeptide repeat proteins (DPRs) generated by repeat-associated non-ATG translation^3– 5^, and haploinsufficiency of C9orf72 due to transcriptional downregulation^1,2^. It remains unclear how each mechanism contributes to neurodegeneration in ALS/FTD^6^.

Recent evidence has implicated dysfunction of nucleocytoplasmic transport (NCT) as an important neurodegenerative pathway in ALS/FTD^7–10^. Indeed, the primary pathology in ALS/FTD affected neurons is nuclear depletion and cytoplasmic aggregation of the normally nuclear TAR DNA-binding protein 43 (TDP-43), suggestive of defective NCT^11–17^. NCT is an essential process in eukaryotic cells by which proteins and RNA transit between the nucleus and cytoplasm through nuclear pore complexes (NPC)^7^; macromolecular channels comprised of numerous nucleoporin proteins (Nups) of varying functions. Small molecules diffuse freely across the NPC, however Ran-GTPase regulates active NCT of cargoes >40kDa^18^, which require nuclear transport receptor (NTR)-mediated trafficking. Importin β-1, one of the most characterized NTRs, binds to transport cargoes in the cytoplasm via the adaptor protein, importin *α*, and facilitates transit across the NPC through interactions with phenyl-alanine (FG)-nucleoporins^19–22^, which line the central channel of the NPC. Active NCT is regulated by the localization and function of Ran-GTPase (the Ran-GTPase cycle)^18^, in which RanGDP in the cytoplasm binds Importin β-1 receptor-cargo complexes and shuttles them into the nucleus. In the nucleus RCC1 (the Ran-GTPase guanine nucleotide exchange factor) stimulates RanGDP to RanGTP exchange, with RanGTP dissociating Importin β-1 and releasing the cargoes. RanGTP associates with export complexes and their cargoes, shuttling them to the cytoplasm where RanGAP (Ran-GTPase activating protein) causes a RanGTP to RanGDP change, releasing the export complexes and cargoes in the cytoplasm. This regulation and maintenance of the Ran-GTPase gradient is critical for defining the directionality and efficiency of NCT^18^.

Evidence of pathological disruption of NCT in ALS/FTD has come from several sources, including the identification of an ALS-causing mutation in *GLE1*, an RNA export factor^23^. Mislocalization of NCT proteins, such as Nups^15,24–28^, NTRs^25,26^ and members of the Ran-GTPase cycle^24,27,29^, are present in disease-affected neurons in both sporadic and *C9orf72*-ALS cases, indicative of NCT dysfunction. Moreover, model systems have demonstrated that ALS-associated species of TDP-43 cause abnormalities in NCT^15,17^, suggesting that mislocalized TDP-43 could not only result from abnormal NCT but also induce further deleterious effects. Crucially, Importin β-1 has been identified as a major determinant of TDP-43 subcellular localization^11^ and disaggregates pathological forms of TDP-43^15,30^.

Intriguingly, both RNA toxicity and DPRs arising from the repeat expansion in *C9orf72* have been shown to disrupt NCT^31–36^. Genetic screens *in vitro*^37^ and *in vivo*^24,38–40^ have identified numerous NCT proteins that act as suppressors or enhancers of G_4_C_2_ repeat expansion RNA- or DPR-mediated phenotypes. These findings also extend to human samples; for example, RanGAP modifies (G_4_C_2_)/DPR phenotypes, binds G_4_C_2_ RNA *in vitro* and colocalizes with (G_4_C_2_) RNA foci in iPSC-derived neurons and postmortem tissue^24^. Moreover, Importin β-1 has been demonstrated to interact directly with arginine-rich DPRs^35,41^. The convergence of (G_4_C_2_) RNA and DPR toxicities onto dysfunctional NCT is striking, however it remains unclear if loss of C9orf72 function also has deleterious effects on NCT.

In *C9orf72*-ALS/FTD, the G_4_C_2_ repeat expansions cause transcriptional downregulation of the *C9orf72* mRNA^1^ leading to reduced levels of C9orf72 protein^29,42–46^. It is therefore critical to establish the function(s) of C9orf72 to understand how downregulation contributes to neurodegeneration. Functionally, C9orf72 exists in a complex with SMCR8 and WDR41^47–50^ and has recently been identified as a GTPase activating protein with multiple targets^51,52^, supporting a role in intracellular trafficking. Studies mainly undertaken *in vitro* have linked C9orf72 with several biological processes including autophagy, endosomal trafficking, lysosomal function, synaptic function, and actin dynamics^42,46–48,50,53–58^. However, *C9orf72* knockout mice do not develop overt neurodegeneration^48,53,59–63^ and primarily show autoimmune phenotypes^60,61,63^, illustrating that *C9orf72* haploinsufficiency alone is insufficient for disease manifestation. Nevertheless, our previous work supports the possibility that maintaining or elevating C9orf72 levels may be protective against neurodegeneration^64^.

We have previously demonstrated that C9orf72 interacts with Ran-GTPase and Importin β-1^29^. Additional findings have also demonstrated that C9orf72 and SMCR8 interact with Importin β-1, NPC proteins and regulators of the Ran-GTPase cycle^46,65,66^. Given the convergence of the other *C9orf72* pathomechanisms onto NCT, here we have investigated how loss of *C9orf72* affects NCT through identifying the subcellular localizations of NCT proteins, including Ran-GTPase and Importin β-1 in *C9orf72* knockout mice. We demonstrate that loss of C9orf72 disrupts the Ran-GTPase gradient both *in vitro* and *in vivo*, and this is associated with defects in NCT. Importantly, we describe neuronal-specific phenotypes in *C9orf72* heterozygous and homozygous knockout mice, identifying compositionally distinct cytoplasmic Importin β-1 granules that variably co-localize with RanGAP. We provide evidence that these granules are derived through budding from the nuclear envelope and are co-labelled with G3BP1 and K63-ubiquitin. Thus, we have established that loss of C9orf72 converges with other *C9orf72* pathomechanisms onto defects in NCT and have identified a neuronal specific function for C9orf72.

## Methods

### Generation of CRISPR-Cas9 C9orf72 knockout HeLa cells

*C9orf72* knockout HeLa cells were generated using a similar strategy as previously described^67^. In brief, guide RNAs (gRNAs) targeting intron 1 (gRNA1: CAACAGCTGGAGATGGCGGT) and exon 2 (gRNA2: GTCCTAGAGTAAGGCACATT) of *C9orf72* were designed using the “optimized CRISPR design” tool (www.crisp.mit.edu). Oligonucleotides with Bsb1 cleavage overhang (Life Technologies) were annealed and cloned into Cas9/puromycin expression vector (PX459 from Addgene #48139). HeLa cells were cultured in Dulbecco’s Modified Eagle’s Medium (DMEM) with high glucose, supplemented with 10% fetal bovine serum (FBS), 100 U/mL penicillin and 100 µg/mL streptomycin at 5% CO_2_ and 37 °C. Genome editing plasmids expressing either gRNA1 or 2 were co-transfected using jetPRIME Transfection Reagent (Polyplus) according to the manufacturer’s protocol. Transfected cells were selected using DMEM supplemented with 3μg/ml puromycin for 24 h. After 96 hours recovery, surviving cells were isolated by clonal dilution and genotyped with polymerase chain reaction (PCR) primers external to the targeted region. Genomic DNA was extracted with QuickExtract (Lucigen) and PCR was performed using Q5® High-Fidelity DNA Polymerase according to the manufacturer’s protocol (Forward primer: ACAGGATTCCACATCTTTGACTT. Reverse primer: TATGTGCTGCGATCCCCATT). *C9orf72* knock out lines were identified using agarose gel electrophoresis and confirmed by Sanger sequencing.

### HeLa cell transfection and chemical treatments

HeLa cells were maintained as above and plated on coverslips pre-coated with poly-d-lysine in 24-well plates (Nunclon). After 24 hours, cells at approximately 85% confluency were transfected using Lipofectamine LTX (Invitrogen), as per the manufacturer’s instructions. The coding sequence of human C9-L was cloned into a pcDNA3.1 plasmid and verified by sequencing. NCT report construct pLVX-EF1alpha-2xGFP:NES-IRES-2xRFP:NLS was a gift from Fred Gage (Addgene plasmid # 71396 ; http://n2t.net/addgene:71396 ; RRID:Addgene_71396)^68^. For chemical treatments, cells were treated with Leptomycin B (10ng/ml) or ethanol for 1 hour. Cells were fixed with 4% paraformaldehyde (PFA) or ice-cold methanol (for Importin β-1 labeling) prior to immunofluorescence labeling.

### Protein extraction and Western blotting

For protein extraction, HeLa cells were grown in 6-well plates, as above, then washed once with ice-cold PBS before lysis with 150µl RIPA buffer (50mM Tris HCl pH7.8, 150mM NaCl, 0.5% sodium deoxycholate, 1% NP40; supplemented with protease inhibitors and EDTA). The cells were harvested, briefly vortexed, and placed on ice for 20 minutes. Lysates were then centrifuged at 20,000xg for 20 minutes at 4ºC and supernatant taken. Protein concentration was determined by bicinchoninic acid (BCA) assay. Sample buffer (187.5mM of Tris-HCl pH 6.8, 6% SDS, 30% glycerol, 0.03% bromophenol blue, 15% β-mercaptoethanol) was added and the samples heated at 95ºC for 5 minutes.

Standard methods for immunoblotting were used. Briefly, homogenates were electrophoresed on 10% (w/v) sodium dodecyl sulphate–polyacrylamide gels and transferred to polyvinylidene fluoride (PVDF) membranes, which were then blocked for 1 hour at ambient temperature in Tris-buffered saline (TBS) containing 5% (w/v) skimmed milk powder. Membranes were incubated overnight at 4°C with primary antibodies diluted in blocking solution as follows: mouse anti-C9orf72 (Genetex, 1:1000), mouse anti-GAPDH (Abcam ab8245, 1:5000), mouse anti-Importin β-1 (Santa Cruz sc-137016, 1:1000) and mouse anti-Ran-GTPase (BD Biosciences 610340, 1:1000). Immunoblots were then washed with TBS containing 0.05% Tween 20 (TBST) and incubated for 1 hour at ambient temperature with either anti-mouse horseradish peroxidase conjugated (NA931, VWR; 1:5000); or anti-rabbit HRP conjugated (NA934, VWR; 1:5000) antibodies diluted in blocking solution. Antibody labelling was visualized using Western Lightning ® Plus ECL (Perkin Elmer) and signal quantified from densitometric scans of autoradiography film (Denville scientific) using ImageJ (NIH).

### C9orf72 knockout mice

*C9orf72* knockout mice on C56BL/6 background were generously provided by Clotilde Lagier-Tourenne (Harvard Medical School) and Don Cleveland (UCSD)^59^. *C9orf72*^*+/-*^ mice were bred to generate *C9orf72*^*+/+*^, *C9orf72*^*+/-*^ and *C9orf72*^*-/-*^ littermates and genotyped as previously described^59^. All procedures were conducted in accordance with the Canadian Council on Animal Care and approved by the University of Toronto Animal Care Committee.

### Primary neuronal cultures

Primary motor neuron cultures from dissociated spinal cords of E13.5 *C9orf72*^*+/+*^ and *C9orf72*^*-/-*^ embryos were prepared as previously described^69^. In brief, cells were plated in Nfeed media (Minimum Essential Media, 1% N3 media, 1.3% horse serum, 0.025% NGF, 0.005% dextrose, 0.0015% sodium bicarbonate; N3 media contains 1mg/ml insulin, 20mg/ml transferrin, 1mg/ml BSA, 20mM putrescine, 15µM selenium, 3µM T3, 2.5µM hydrocortisone and 320µM progesterone), supplemented with 1% penicillin-streptomycin and 1% FBS at a density of 250,000 cells per well in 24 well plates (Nunclon) on coverslips precoated with poly-d-lysine and Matrigel (VWR) diluted 1:200 in Minimum Essential Media (HyClone). After 4 days *in vitro* (DIV), media was replaced with Nfeed containing 5’-fluoro-2-deoxyuridine (FDU) to limit the growth of mitotic cells. After 6DIV, FDU-containing Nfeed was removed and replaced with Nfeed. Half media changes were performed every 2-3 days. After 21 days *in vitro* (DIV) cells were fixed in 4% PFA or ice-cold methanol prior to immunofluorescence labeling.

Primary cortical neurons were isolated from brains of E15 *C9orf72*^*+/+*^ and *C9orf72*^*-/-*^ mouse embryos. Following trypsin digestion and dissociation, cells were plated (250,000 cells per well in a 24 well plate) in plating media (DMEM, 1% penicillin-streptomycin, Glutamax (Gibco), 10% horse serum) onto coverslips precoated overnight with poly-d-lysine (4mg/ml) in sodium borate buffer. After 2 hours, plating media was replaced with Neurobasal Media™ (Thermo Fisher Scientific) supplemented with 1% penicillin-streptomycin, GlutaMAX and B27 (Thermo Fisher Scientific). After 4DIV, FDU was added and removed at 6DIV. Half media changes were performed every 2-3 days and cells were fixed in 4% PFA or ice-cold methanol after 7DIV prior to immunofluorescence labeling.

### Immunofluorescence labeling of HeLa cells and primary neuronal cultures

After fixation, as above, cells on coverslips were permeabilized with 0.3% Triton X-100 in phosphate buffered saline (PBS) for 20 minutes. Cells were then blocked and permeabilized in 10% donkey serum in PBS containing 0.3% Triton-X100 for 1 hour at room temperature, then incubated with primary antibodies diluted in blocking solution overnight at 4ºC. Following 3×5 minute PBS washes, the appropriate Alexa Fluor™ (Invitrogen) secondary antibodies diluted in PBS were applied to coverslips for 1 hour at room temperature and protected from light. After 3×5 minute PBS washes, coverslips were mounted on slides with *ProLong*® *Gold Antifade Reagent* containing DAPI. Primary antibodies used for immunofluorescence of HeLa cells and/or primary neurons were: mouse anti-Ataxin-2 (BD Biosciences 611378, 1:250), mouse anti-beta-III-tubulin (Covance MMS-435P, 1:1000), mouse anti-CRM1 (Abcam ab191081, 1:250), goat anti-Importin β-1 (Santa Cruz sc-1863, 1:50), mouse anti-Importin β-1 (Santa Cruz sc-137016, 1:50), mouse anti-KPNA (BD Biosciences 610485, 1:200), goat anti-Lamin B (Santa Cruz sc-6217, 1:200), rabbit anti-MAP2 (Millipore AB5622, 1:1000), rabbit anti-Nup98 (Santa Cruz sc-74553, 1:50), rabbit anti-RanBP1 (Abcam ab97659, 1:200), rabbit anti-RanBP2 (Abcam ab ab64276, 1:200), rabbit anti-RanGAP (Santa Cruz sc-25630, 1:250), mouse anti-Ran-GTPase (BD Biosciences 610340, 1:500), goat anti-RCC1 (Santa Cruz sc-1161, 1:50), mouse anti-TNPO (Abcam ab10303, 1:200).

### Harvesting and processing of mouse tissue

Adult mice at 2 or 6 months of age were anesthetized with an intraperitoneal injection of ketamine/xylazine (100/10 mg/kg body weight) then transcardially perfused with PBS, followed by 10% neutral buffered formalin (Sigma). Brain and spinal cords were removed, and immersion fixed for 1 week in formalin, followed by storage in 70% ethanol. Tissues were paraffin embedded and 6µm sections cut using a microtome onto charged slides. For neonatal mice, tissues were dissected and immersion fixed for 1 week in formalin, then stored in 70% ethanol.

### Immunofluorescence on mouse formalin-fixed paraffin-embedded tissue

Sections on slides were heated at 60ºC for 25 minutes for deparaffinization and rehydrated through a series of washes in graded ethanol and finally in water, as previously described^29,57,64,70^. Antigen retrieval was performed by pretreating sections with TE9 buffer (10mM Trizma base, 1mM EDTA, 0.2% Tween 20 [pH 9]), at 110 °C in a pressure cooker for 15 minutes, followed by cooling for 20 minutes and washing in running tap water for 20 minutes. After a 5 minute wash in Tris-buffered saline (TBS) containing 0.05% Tween®-20 (TBS-T), tissue was permeabilized by incubation in TBS containing 0.3% Triton-X100 for 20 minutes at room temperature, followed by a 5 minute wash in TBS-T. After 1 hour incubation with blocking solution (0.3% Triton-X100, 3% bovine serum albumin, 5% donkey or goat serum), slides were incubated overnight at 4ºC with primary antibodies diluted in antibody diluent (Dako). Following 3×10 minute TBS-T washes, appropriate Alexa Fluor™ secondary antibodies diluted in Dako antibody diluent were applied for 1 hour at room temperature. Following 3×10 minute TBS-T washes, slides were mounted with coverslips using *ProLong*® *Gold Antifade Reagent* containing DAPI. Primary antibodies used for immunofluorescence of mouse tissue were: mouse anti-Ataxin-2 (BD Biosciences 611378, 1:250), rabbit anti-Caprin1 (Proteintech, 15112-1-AP, 1:500), rabbit anti-G3BP1 (Proteintech 13057-2-AP, 1:500), goat anti-Importin β-1 (Santa Cruz sc-1863, 1:50), mouse anti-Importin β-1 (Santa Cruz sc-137016, 1:50), rabbit anti-Importin β-1 (Santa Cruz sc-11367, 1:50), goat anti-Lamin B (Santa Cruz sc-6217, 1:200), rabbit anti-LC3 (Abcam, ab128025, 1:200), mouse anti-mAb414 (Abcam, ab24609, 1:100), rabbit anti-MAP2 (Millipore AB5622, 1:1000), mouse anti-NeuN (Millipore, MAB377, 1:500), rabbit anti-Nup205 (Novus Bio, NBP1-91247, 1:200), rabbit anti-p62 (MBL, PM045, 1:250), rabbit anti-POM121 (ThermoFisher PA5-36498, 1:250), rabbit anti-RanBP2 (Abcam ab63276, 1:200), rabbit anti-RanGAP (Santa Cruz sc-25630, 1:200), mouse anti-Ran-GTPase (BD Biosciences 610340, 1:250), mouse anti-SMI-32 (Biolegend, SMI32P, 1:500), rabbit anti-TDP-43 (Proteintech 10782-2-AP, 1:2000), rabbit anti-Ubc9 (Abcam, ab33044, 1:200), mouse anti-Ubiquitin (Millipore, MAB1510, 1:250), rabbit anti-Ubiquitin, Lys48(K48)-specific (Millipore, 05-1307, 1:50), rabbit anti-Ubiquitin, Lys63(K63)-specific (Millipore, 05-1308, 1:50).

### Microscopy and Quantification of Ran-GTPase, GFP, TDP-43 N:C ratios

To quantify nuclear:cytoplasmic (N:C) ratios of Ran-GTPase, GFP or TDP-43, z-stack images were taken at 0.5µm intervals using the x63 objective of a Leica DM6000B microscope with images acquired using Volocity software (PerkinElmer®). The nuclear and total cell body boundaries were manually drawn using Volocity software, and fluorescence values obtained. For higher resolution imaging, images were acquired using the 63x objective on a spinning disk confocal microscope and Z-stack images taken with an interval size of 0.13µm. Volocity software was used to deconvolve images and produce 3D renderings.

### Quantification of cytoplasmic Importin β-1 granules and colabeling

The investigator was blind to genotype and age group. Z-stack images at 0.5µm intervals were acquired using the x63 objective of a Leica DM6000B microscope from dentate gyrus (DG), CA3, CA1, cortex and cerebellar Purkinje cells. Total number of cells containing Importin β-1 or RanGAP granules were scored and expressed by the area of the region examined. In the lumbar spinal cord, the total number of cytoplasmic Importin β-1 granules was expressed per motor neuron examined. Due to different morphologies of Importin β-1 granules in spinal cord motor neurons, granules were also scored as small (S-granules) or large (L-granules). To calculate colocalization percentages, Importin β-1 granules were scored for colabeling with RanGAP, and vice versa. A similar strategy was used to calculate colocalization of Importin β-1 granules with G3BP1 and ubiquitin. ImageJ (NIH) was used to measure granule area.

## STAR Methods

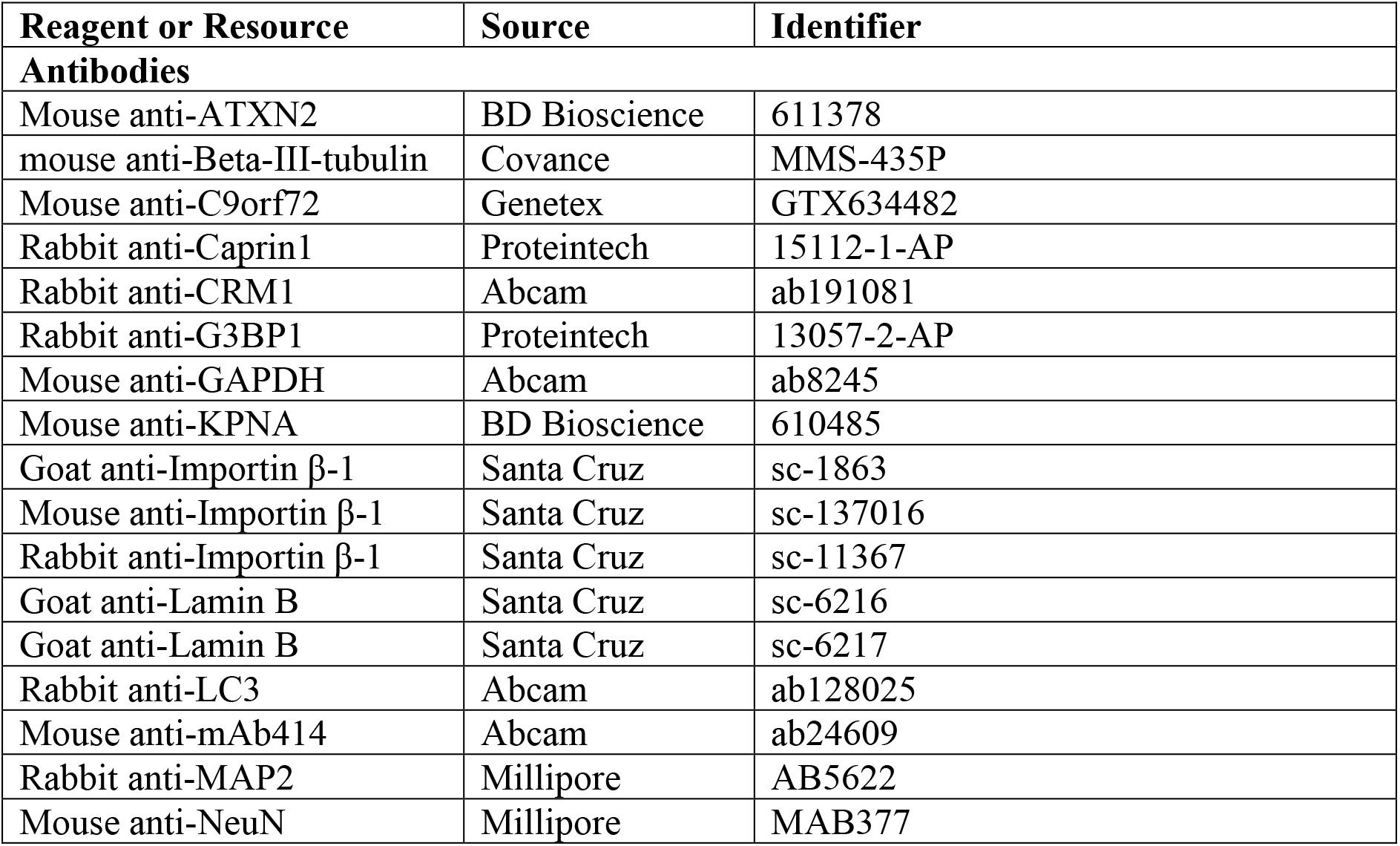
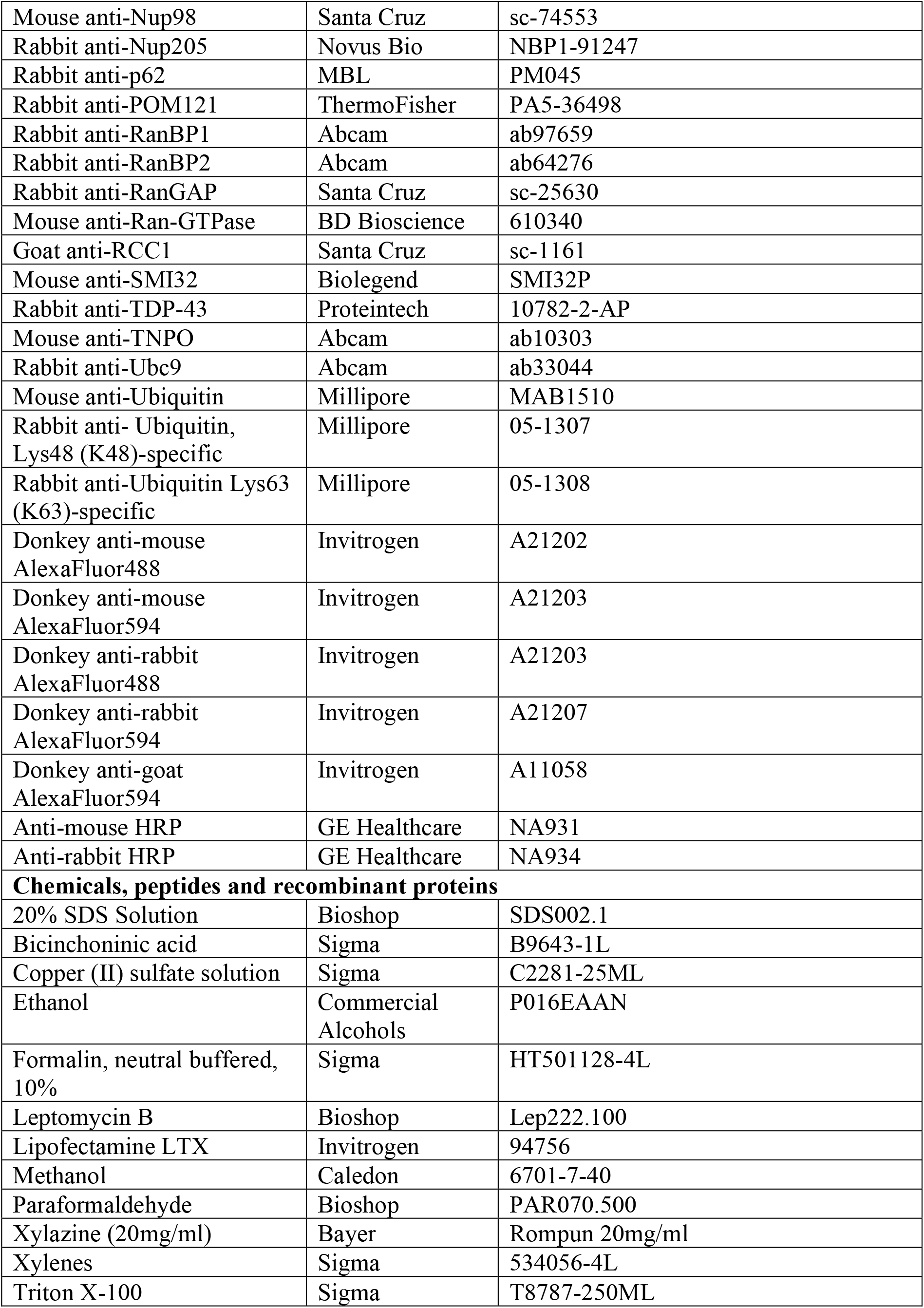
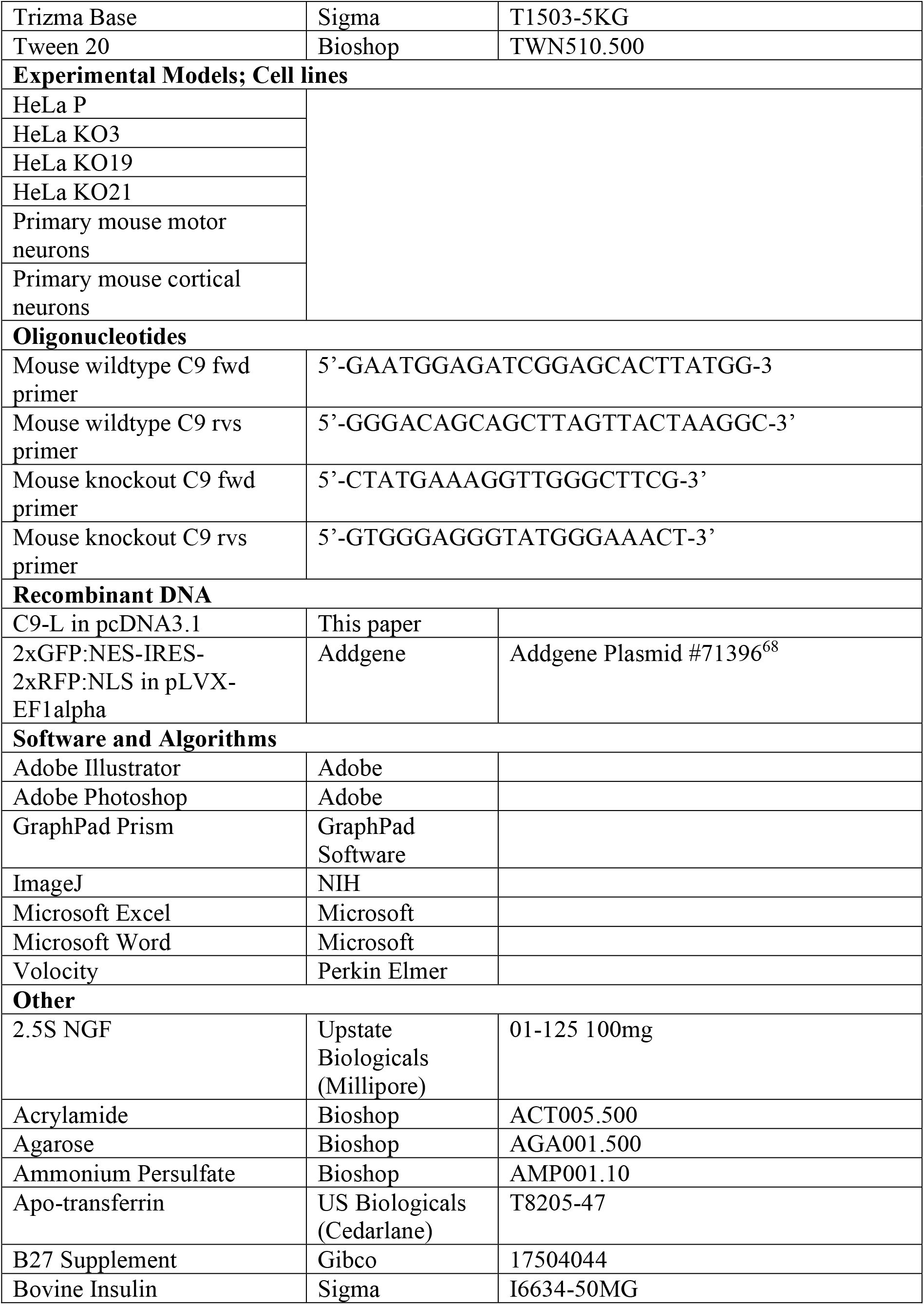
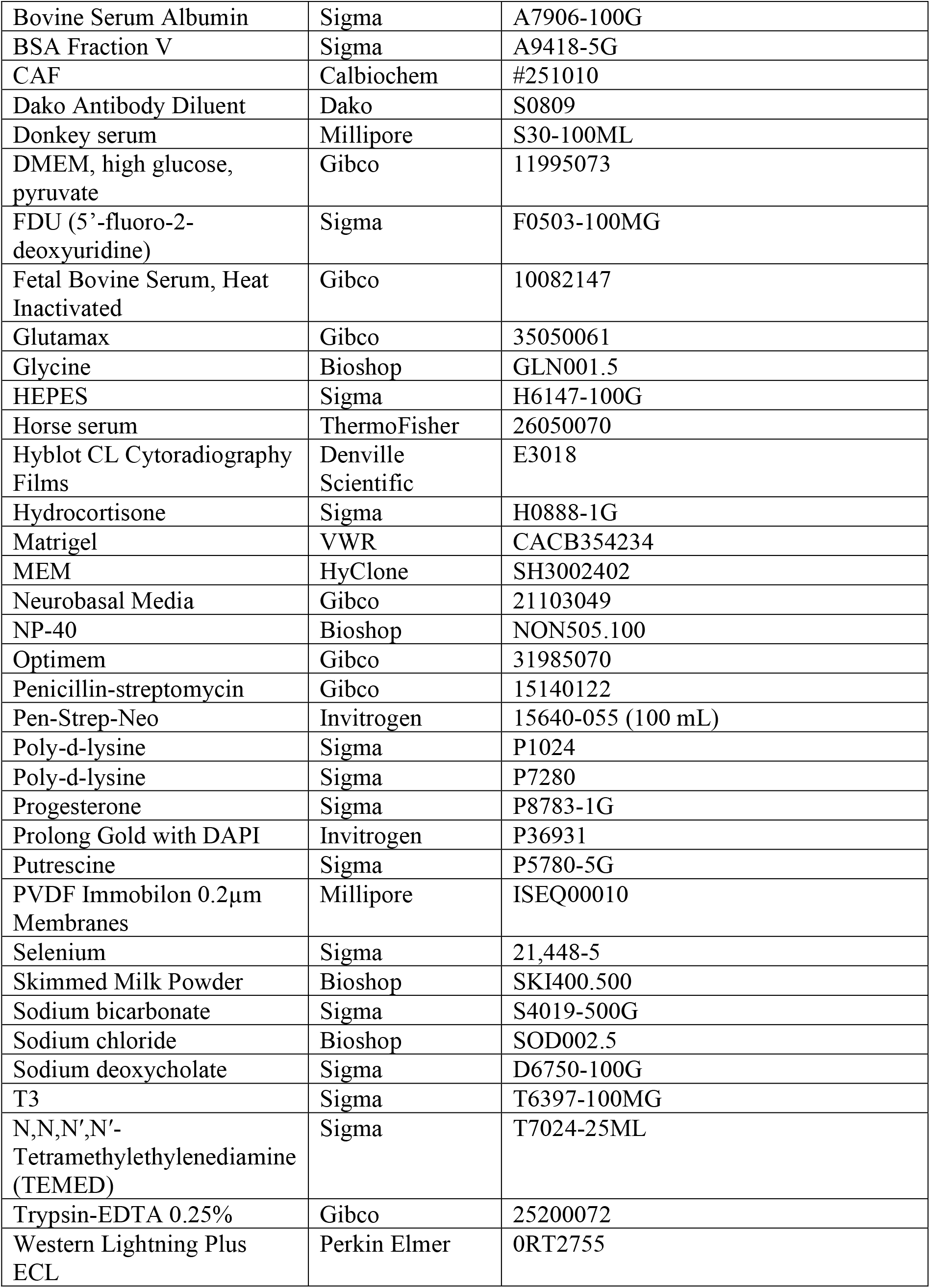

## Results

### Loss of *C9orf72* disrupts the Ran-GTPase gradient and nucleocytoplasmic transport in HeLa cells

To investigate the role of C9orf72 in NCT, knockout HeLa cells were generated using CRISPR-Cas9 with guide RNAs designed to induce deletions and frameshifts in the endogenous *C9orf72* gene, thereby disrupting its protein production. Genomic DNA analysis confirmed deletions in the *C9orf72* gene (**Figure S1A**). In humans there are two protein isoforms of C9orf72: C9-S (approximately 23kDa) and the predominant isoform C9-L (approximately 52kDa), which is the more widely characterized protein^1,2,29,71,72^. Using a recently validated antibody^67^, C9-L protein was detectable in the parental cell line but not in three independently generated knockout lines (**Figure S1B**). Although Western blot analysis was performed using a relatively high amount of protein (120µg), we did not detect C9-S in parental or knockout lines; thus, if C9-S is expressed in parental lines its expression is extremely low. As we have previously demonstrated that C9orf72 interacts with Ran-GTPase and Importin β-1^29^, their protein levels were quantified by Western blot, with no differences observed in *C9orf72* knockout lines compared to the parental line, suggesting that loss of C9orf72 does not affect expression of Ran-GTPase or Importin β-1 (**Figure S1C**).

The nuclear:cytoplasmic (N:C) ratio of Ran-GTPase (Ran gradient) can be used to assay its function^24^. This was quantified by measuring the nuclear versus cytoplasmic fluorescence intensities of Ran-GTPase labeling in control and *C9orf72* knockout HeLa cells (**Figure 1A, Figure S1D**). *C9orf72* knockout lines showed a significant decrease in Ran-GTPase N:C ratio (**Figure 1B**). Similar changes have been previously reported in iPSC-derived neurons from *C9orf72* patients^24,73^. To determine whether the altered Ran-GTPase N:C ratio caused by loss of C9orf72 affected NCT, parental and *C9orf72* knockout HeLa cells were transfected with a reporter construct (pLVX-EF1alpha-2xGFP:NES-IRES-2xRFP:NLS) that was previously used to identify age-dependent^68^ and Tau-induced NCT abnormalities^74^. In parental cells, GFP (2xGFP:NES; nuclear export signal) localized to the cytoplasm and RFP (2xRFP:NLS; nuclear localization signal) to the nucleus (**Figure 1C, Figure S1E**). Therefore, the N:C ratio of these reporter proteins can be used to measure the efficiency of NCT. To confirm the validity of the reporter distribution, cells expressing pLVX-EF1alpha-2xGFP:NES-IRES-2xRFP:NLS were treated with Leptomycin B, an inhibitor of nuclear export^14,68^, which caused accumulation of GFP in the nucleus (**Figure S1E and S1F**). In the *C9orf72* knockout lines, the N:C ratio of the 2xGFP:NES was significantly increased compared to the parental line (**Figures 1C and 1D, Figure S1G**), indicating that loss of C9orf72 not only caused mislocalization of Ran-GTPase but also dysfunction of NCT.

**Figure 1:**
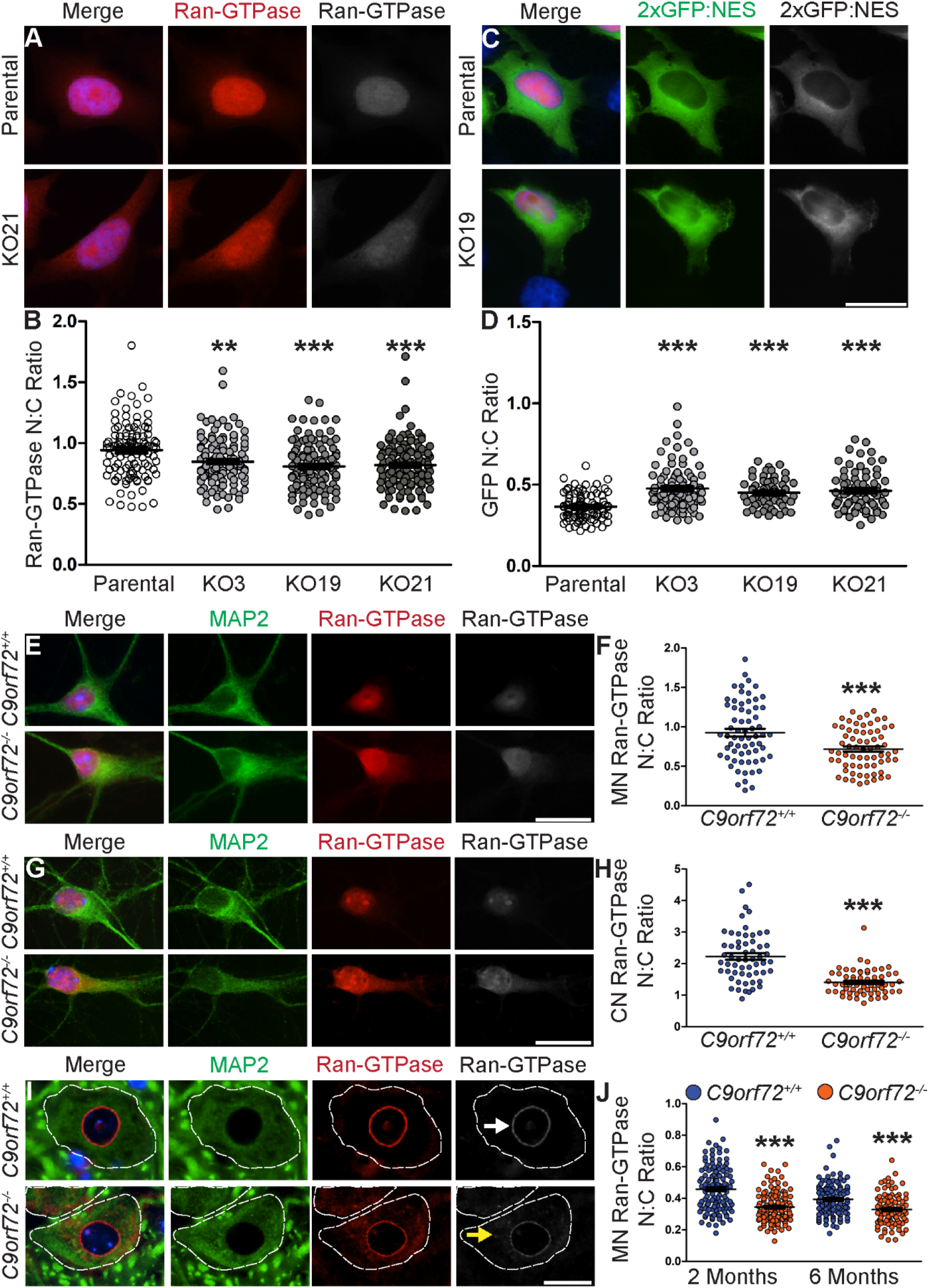
Loss of C9orf72 disrupts Ran-GTPase localization in HeLa cells, cultured primary neurons and spinal motor neurons in vivo. **(A)** Representative images of immunofluorescence labeling of Ran-GTPase in parental and *C9orf72* knockout (KO21) HeLa cells. Note increased cytosolic Ran-GTPase in KO21 compared to parental control. **(B)** Quantification of the Ran-GTPase nuclear:cytoplasmic (N:C) ratio revealed a significant decrease in *C9orf72* knockout lines (KO3, KO19, KO21) compared to parental control (parental= 0.94±0.02, KO3=0.85±0.02, KO19=0.81±0.02, KO21=0.82±0.02 ; n≥100 cells per line, p<0.01=**, p<0.001=***). **(C)** HeLa cells were transfected with pLVX-EF1alpha-2xGFP:NES-IRES-2xRFP:NLS reporter construct, with 2xGFP:NES (nuclear export signal-green) found predominantly in the cytoplasm in the parental line. Note increased nuclear localization of GFP in the representative knockout cell line (KO19). **(D)** Quantification of the GFP:NES reporter N:C demonstrated a significant increase in *C9orf72* knockout cells compared to parental control, indicative of NCT dysfunction (parental=0.36±0.01, KO3=0.48±0.01, KO19=0.45±.01, KO21=0.46±0.01; n≥60 cells per line, p<0.001=***). **(E-J)** Double immunofluorescence was performed for neuronal marker MAP2 (green) and Ran-GTPase (red), followed by Ran-GTPase N:C ratio quantification, on primary motor neurons **(E, F)** and primary cortical neurons **(G, H)** established from *C9orf72*^*+/+*^ and *C9orf72*^*-/-*^ mouse embryo littermates, at 21DIV and 7DIV, respectively, and *C9orf72*^*+/+*^ and *C9orf72*^*-/-*^ spinal cord motor neurons at 2 and 6 months of age **(I, J)**. In spinal cord motor neurons note localization of Ran-GTPase to the nuclear envelope (white arrow) and increased cytoplasmic localization of Ran-GTPase in *C9orf72*^*-/-*^ compared to *C9orf72*^*+/+*^ motor neurons (yellow arrow). Quantification of the Ran-GTPase N:C ratio revealed a significant decrease in *C9orf72*^*-/-*^ primary motor neurons (**F**; *C9orf72*^*+/+*^=0.93±0.05, *C9orf72*^*-/-*^=0.72±0.03; n≥50 neurons per genotype, p<0.001=***), primary cortical neurons (**H**; *C9orf72*^*+/+*^=2.23±0.01, *C9orf72*^*-/-*^=1.41±0.05; n≥55 neurons per genotype, p<0.001=***) and spinal cord motor neurons (**J**; 2 months: *C9orf72*^*+/+*^=0.45±0.13, *C9orf72*^*-/-*^=0.34±0.01; n≥120 motor neurons per genotype, p<0.001=***; 6 months: *C9orf72*^*+/+*^=0.439±0.01, *C9orf72*^*-/-*^=0.33±0.01, n≥90 motor neurons per genotype, p<0.001=***) compared to *C9orf72*^*+/+*^ neurons. Motor neuron cell bodies are denoted by white dotted line. DAPI nuclear stain (blue). Unpaired two-tailed t-tests were used for statistical analysis. Data are mean±SEM. Scale bars = 20µm. CN=cortical neuron, MN=motor neuron, NES=nuclear export signal.

NCT dysfunction is frequently characterized by the mislocalization of proteins of the Ran-GTPase cycle, import/export receptors or NPC proteins^15,24,25,28,35,38–40^. Given the mislocalization of Ran-GTPase and the associated NCT dysfunction we observed in *C9orf72* knockout HeLa cells, we next examined the subcellular localization of proteins involved in these processes. Immunofluorescence labelling of proteins involved in the Ran-GTPase cycle (RanBP1, RanBP2, RanGAP, RCC1; **Figures S2A-D**), nuclear import and export (KPNA1, TNPO, CRM1, Importin β-1; **Figures S2E-H**), NPC and nuclear envelope (Nup98, **Figure S2I;** Lamin B; **Figure S2J**), and ataxin-2 (**Figure S2K**) revealed similar localization patterns of each protein in parental and *C9orf72* knockout lines. These data suggest that in HeLa cells, loss of C9orf72 disrupts Ran-GTPase localization and NCT function without affecting the subcellular localization of other proteins involved in NCT.

### The Ran-GTPase gradient is disrupted in neurons lacking C9orf72

To determine if loss of C9orf72 induced similar effects on Ran-GTPase distribution in the neuronal subtypes affected in ALS/FTD, primary motor (**Figure 1E and 1F**) and cortical (**Figure 1G and 1H**) neurons were established from *C9orf72*^*+/+*^ and *C9orf72*^*-/-*^ mouse embryos and Ran-GTPase N:C assessed by measuring the respective N:C fluorescence intensities. In motor neurons at 21 days *in vitro* (DIV), the Ran-GTPase N:C ratio was significantly decreased in *C9orf72*^*-/-*^ neurons compared to *C9orf72*^*+/+*^ neurons (**Figure 1F**). Similar findings were evident in primary cortical neurons at 7 DIV (**Figure 1H**). We next examined whether loss of C9orf72 induced Ran-GTPase mislocalization in neurons of *C9orf72*^*-/-*^ mice *in vivo* (**Figures 1I and 1J**). Due to the inflammatory and autoimmune phenotypes that develop in *C9orf72*^*-/-*^ mice^60,61,63,75–77^, we selected 2 month and 6 month timepoints to minimize the effects of these confounds. The Ran-GTPase N:C was assessed in motor neurons of the lumbar spinal cord, chosen due to their vulnerability in ALS. In contrast to the diffuse nuclear and cytoplasmic localization of Ran-GTPase in cultured cells (**Figures 1A, 1E, 1G, Figure S1D**), Ran-GTPase was primarily localized to the nuclear envelope in *C9orf72*^*+/+*^ spinal motor neurons (**Figure 1I**, white arrow) and exhibited increased localization to the cytoplasm in *C9orf72*^*-/-*^ knockout motor neurons (**Figure 1I**, yellow arrow). The Ran-GTPase N:C ratio in motor neurons was significantly reduced in *C9orf72*^*-/-*^ mice compared to *C9orf72*^*+/+*^ mice at 2 months of age, with similar effects at 6 months of age (**Figure 1J**). Thus, our analysis demonstrates that loss of C9orf72 in neurons *in vitro* and *in vivo* causes cytoplasmic mislocalization of Ran-GTPase and that this is not changed between 2 and 6 months of age.

### Increased abundance of cytoplasmic Importin β-1 granules in *C9orf72* knockout motor neurons

Based on our previous finding of an interaction between C9orf72 and Importin β-1 in cultured cells^29^, we examined the subcellular localization of Importin β-1 in motor neurons of the lumbar spinal cord of *C9orf72*^*+/+*^ and *C9orf72* knockout mice. Although Importin β-1 was not mislocalized in *C9orf72* knockout HeLa cells or primary motor neurons (**Figure S2L**), we speculated that because NCT function declines with age^68^, the combination of age and reduced C9orf72 would manifest phenotypes not seen in the lifespan of cultured cells. Importin β-1 antibody labelling revealed diffuse localization within the nucleus and strong labeling of the nuclear envelope in spinal motor neurons of *C9orf72*^*+/+*^, *C9orf72*^*+/-*^ and *C9orf72*^*-/-*^ mice (**Figure 2A**). Unexpectedly, Importin β-1 antibody also labelled cytoplasmic granules (**Figure 2A**), present in both the soma and axons of motor neurons **(Figure 2B)**. Cytoplasmic Importin β-1 granules were spherical in shape and two clear sizes could be distinguished (see boxes in **Figures 2A-C**): small granules (S-granules, white arrows), with an average size of 0.15µm^2^±0.01µm^2^ (p<0.01), and large-granules (L-granules; yellow arrows) with an average size of 0.30µm^2^±0.02µm^2^ (p<0.01) (**Figures 2C and 2D**). These sizes were consistent between genotypes.

**Figure 2:**
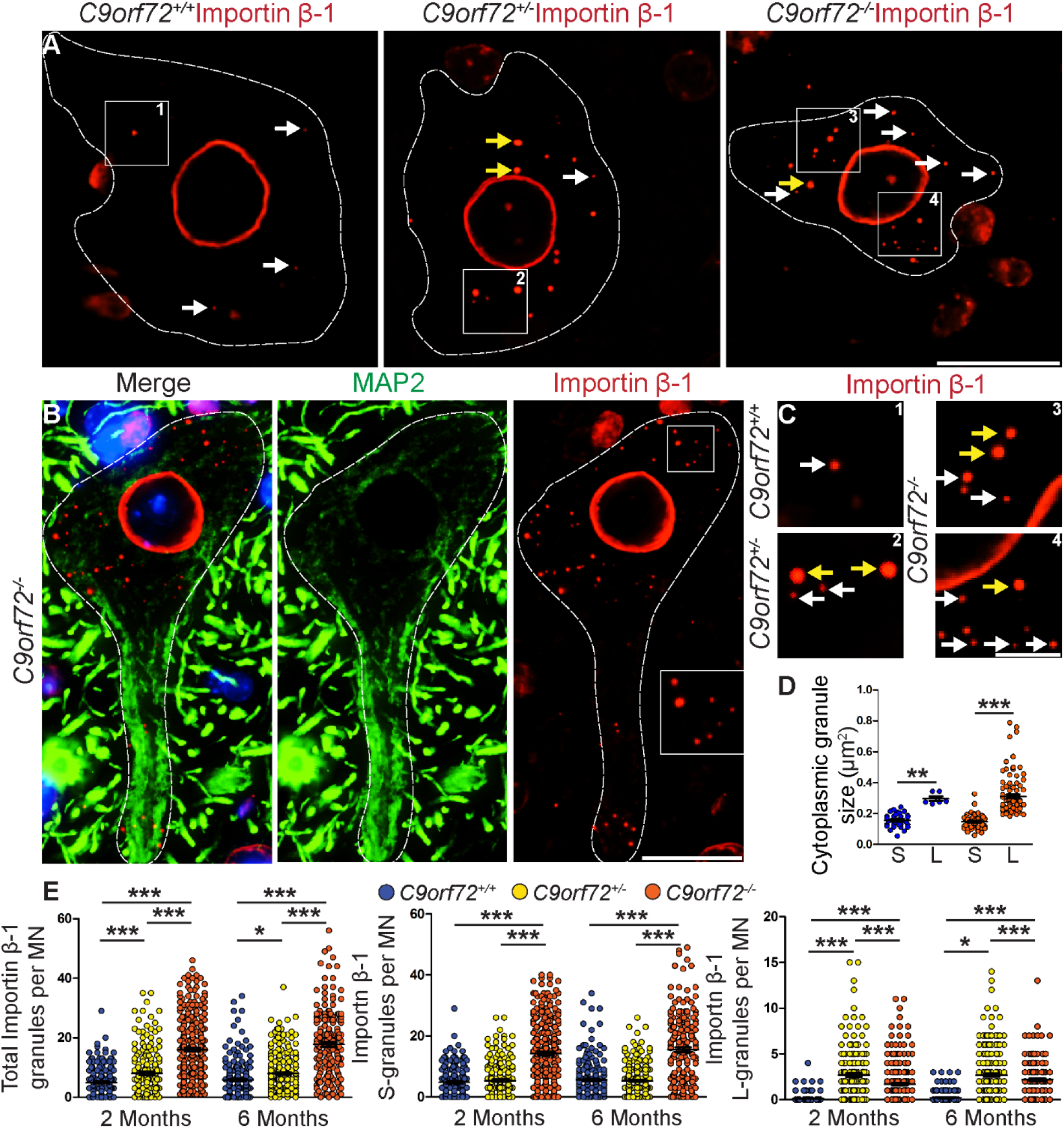
Loss of C9orf72 causes increased abundance of cytoplasmic Importin β-1 granules in spinal motor neurons. **(A)** Immunofluorescence labelling of Importin β-1 in spinal motor neurons of *C9orf72*^*+/+*^, *C9orf72*^*+/-*^ and *C9orf72*^*-/-*^ mice. Note strong labeling of nuclear envelope and presence of cytoplasmic granules that appear to have increased abundance in *C9orf72* knockout neurons. Boxed. **(B)** Double immunofluorescence labelling for neuronal marker MAP2 (green) and Importin β-1 (red) demonstrates that both S- and L-Importin β-1 granules were present in soma and axons. DAPI nuclear stain (blue). **(C and D)** Boxed areas (1-4, from **A**) magnified in **(C)** indicate different sizes of Importin β-1 granules. Measurement of granules **(D)** revealed two populations: S-granules of approximately 0.15µm^2^±0.01µm^2^ (indicated with white arrows) and L-granules of approximately 0.31µm^2^±0.01 µm^2^ (indicated with yellow arrows). **(E)** Quantification of total cytoplasmic Importin β-1 granules (left panel), Importin β-1 S-granules (middle panel) and Importin β-1 L-granules (right panel) in motor neurons of all genotypes at 2 and 6 months of age. Per genotype at least 224 and 196 motor neurons were counted at 2 and 6 months of age, respectively. For quantification n=3-5 mice per genotype were used. Data are mean±SEM. Motor neuron cell bodies are denoted by white dotted line. One-way ANOVA with Bonferroni post-hoc testing was used for statistical analysis. p<0.05=*, p<0.001=***. Scale bars A, D = 20µm. Scale bar B = 5µm. MN = motor neuron.

Qualitatively, it appeared that there were increased numbers of Importin β-1 granules in *C9orf72*^*+/-*^ and *C9orf72*^*-/-*^ motor neurons compared to *C9orf72*^*+/+*^ motor neurons (**Figure 2A**). Quantification confirmed that at 2 months of age there were on average 4.94±0.29 (range 0-29) total Importin β-1 granules in the cytoplasm of *C9orf72*^*+/+*^ motor neurons (n=242), 8.05±0.47 (range 0-35) in *C9orf72*^*+/-*^ motor neurons (n=224) and 15.98±0.57 (range 1-46) in *C9orf72*^*-/-*^ motor neurons (n=320) (**Figure 2E, Table 1**). Similar numbers were obtained in mice at 6 months of age, showing that there was no age-dependent increase in the number of Importin β-1 granules (**Figure 2E, Table 1**). Separate quantification of S-granules and L-granules revealed that there were largely equivalent numbers of Importin β-1 S-granules in *C9orf72*^*+/+*^ and *C9orf72*^*+/-*^ motor neurons and an approximately 3-fold increase in *C9orf72*^*-/-*^ motor neurons (**Figure 2E, Table 1**). Importin β-1 L-granules were rarely observed in *C9orf72*^*+/+*^ motor neurons, however there was a 25-fold increase of Importin β-1 L-granules in *C9orf72*^*+/-*^ motor neurons and a 15-fold increase in *C9orf72*^*-/-*^ motor neurons at 2 months of age. Similar results were obtained at 6 months of age (**Figure 2E, Table 1**).

**Table 1.** Quantification of cytoplasmic Importin β-1 granules in motor neurons of *C9orf72*^*+/+*^, *C9orf72*^*+/-*^ and *C9orf72*^*-/-*^ mice at 2 and 6 months of age. (n=number of MNs examined). MN=motor neuron.

| **2 Months of Age** | *C9orf72*<sup>+/+</sup><br>(n=242) | *C9orf72*<sup>+/-</sup><br>(n=224) | *C9orf72*<sup>-/-</sup><br>(n=320) |
| --- | --- | --- | --- |
| Average Total Importin β-1 granules per MN | 4.94±0.29<br>(range: 0-29) | 8.05±0.47<br>(range: 0-35) | 15.98±0.57<br>(range: 1-46) |
| Average Importin β-1 S-granules per MN | 4.83±0.29<br>(range: 0-29) | 5.34±0.36<br>(range: 0-26) | 14.28±0.58<br>(range: 0-40) |
| Average Importin β-1 L-granules per MN | 0.11±0.03<br>(range: 0-4) | 2.71±0.18<br>(range: 0-15) | 1.70±0.11<br>(range: 0-11) |
| **6 Months of Age** | *C9orf72*<sup>+/+</sup><br>(n=211) | *C9orf72*<sup>+/-</sup><br>(n=303) | *C9orf72*<sup>-/-</sup><br>(n=196) |
| Average Total Importin β-1 granules per MN | 5.81±0.42<br>(range: 0-34) | 7.95±0.35<br>(range: 0-37) | 17.72±0.79<br>(range: 0-56) |
| Average Importin β-1 S-granules per MN | 5.60±0.41<br>(range: 0-34) | 5.27±0.27<br>(range: 0-26) | 15.54±0.80<br>(range: 0-49) |
| Average Importin β-1 L-granules per MN | 0.20±0.04<br>(range: 0-3) | 2.68±0.14<br>(range: 0-14) | 2.18±0.14<br>(range: 0-13) |

Collectively, we have identified two types of Importin β-1 granules in the cytoplasm of mouse spinal motor neurons, S-granules and L-granules. A 50% reduction of C9orf72 increases the abundance of Importin β-1 L-granules, and complete loss of C9orf72 increases the abundance of both Importin β-1 S-granules and L-granules. It is notable that these granules were not observed in HeLa cells or primary motor neurons (**Figures S2H and S2L**, respectively).

### Loss of C9orf72 dysregulates cytoplasmic NPC proteins

To investigate whether loss of C9orf72 caused widespread mislocalization of NCT components, we performed immunostaining against mAb414 (recognizes FxFG repeat sequences in multiple Nups), RanGAP (a regulator of the Ran-GTPase cycle, responsible for hydrolysis of RanGTP to RanGDP), Ubc9 (interacting-partner of RanGAP involved in regulation of the Ran-GTPase cycle and dissociation of nuclear export complexes^78,79^) and RanBP2 (a regulator of the Ran-GTPase cycle, also known as Nup358). FG-Nups and RanGAP have been implicated in *C9orf72* gain-of-function pathways^24,31,34,38,40,80,81^ and RanBP2 has been shown to interact with SMCR8^65^. RanBP2 also forms a complex with RanGAP and Ubc9 (the RanBP2 complex)^78,79,82^. MAb414 (**Figure 3A**), RanGAP (**Figure 3B**) and Ubc9 (**Figure 3C**) labelled the nuclear envelope and RanBP2 (**Figure 3D**) exhibited diffuse nuclear staining in spinal cord motor neurons of all three genotypes. Importantly, mAb414, RanGAP, Ubc9 and RanBP2 co-localized with cytoplasmic S- and L-Importin β-1 granules in *C9orf72*^*+/+*^ motor neurons. These granules had the appearance of annulate lamellae pore complexes (ALPCs)^83,84^. Supporting this there was no co-labeling of S- or L-Importin β-1 granules with POM121, a transmembrane nucleoporin that does not co-associate with ALPCs (**Figure S3A**)^85^. Interestingly, although there was an increased abundance of Importin β-1 S and L granules in *C9orf72*^*+/-*^ and *C9orf72*^*-/-*^ motor neurons, there was a significant reduction in co-labelling with mAb414, RanGAP, Ubc9 and RanBP2 (**Figures 3A-D**). Of note, there was no labeling of Importin β-1 S and L granules with other proteins associated with NCT and/or ALS (POM121, Lamin B, Nup205, Ataxin-2) (**Figures S3B-E**). Moreover, the N:C ratio of TDP-43 in motor neurons of *C9orf72*^*-/-*^ mice compared to *C9orf72*^*+/+*^ mice was unchanged (**Figures S3F and S3G**).

**Figure 3:**
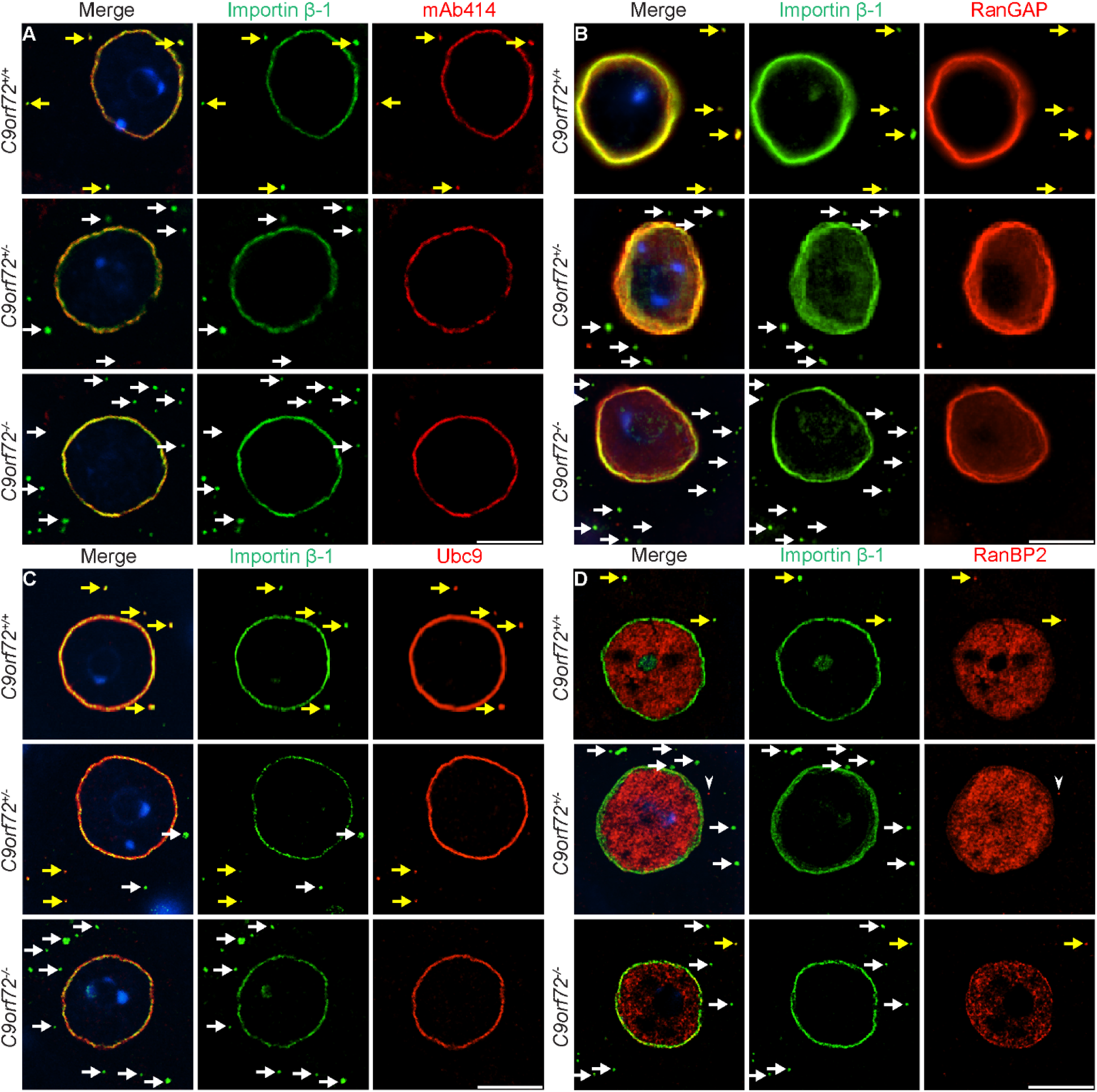
*Loss of C9orf72 disrupts the co-localization of FG-Nups and RanBP2 complex proteins with Importin* β-1 granules in spinal motor neurons. **(A-D)** Double immunofluorescence labelling of Importin β-1 (green) in *C9orf72*^*+/+*^, *C9orf72*^*+/-*^ and *C9orf72*^*-/-*^ motor neurons with mAb414 (**A**, red), RanGAP (**B**, red), Ubc9 (**C**, red) and RanBP2 (**D**, red). Note co-localization of mAb414, RanGAP, Ubc9 and RanBP2 in *C9orf72*^*+/+*^motor neurons (indicated with yellow arrows) and increased abundance of Importin β-1 granules and loss of co-localizations in *C9orf72*^*+/-*^ and *C9orf72*^*-/-*^ motor neurons (indicated by white arrows). Scale bars = 10µm.

These data demonstrate that FG-Nups and RanBP2 complex proteins co-localize with Importin β-1 S- and L-granules in *C9orf72*^*+/+*^ motor neurons. This association is lost in *C9orf72*^*+/-*^ and *C9orf72*^*-/-*^ motor neurons, with the abundance of both Importin β-1 S- and L-granules increased.

### Cytoplasmic Importin β-1 granules are present in cortical and hippocampal neurons of *C9orf72*^*+/-*^ *and C9orf72*^*-/-*^ mice, and colocalize with RanGAP

Based on the presence of cytoplasmic Importin β-1 granules in motor neurons, we examined whether Importin β-1 granules were present in other neuron lineages in the central nervous system. Labeling of *C9orf72*^*+/+*^, *C9orf72*^*+/-*^ and *C9orf72*^*-/-*^ mouse brain with Importin β-1 antibody revealed localization to the nucleus and nuclear envelope in neurons of all genotypes (**Figures 4A and 4B**). Cytoplasmic Importin β-1 granules were observed in the hippocampus (**Figure 4A**), cortex (**Figure 4B**), olfactory bulb (data not shown) and brainstem (data not shown) but not cerebellar neurons (**Figure S4A**) *C9orf72*^*+/-*^ and *C9orf72*^*-/-*^ mice, and were absent in *C9orf72*^*+/+*^ mice (**Figures 4A and 4B**). NeuN co-labelling confirmed that Importin β-1 granules were present in neurons (**Figure S4B**). Importin β-1 granules in *C9orf72*^*-/-*^ cortical and hippocampal neurons were spherical in shape and larger than L-granules in motor neurons (0.44µm^2^±0.02µm^2^, versus L-granules: 0.30µm^2^±0.02µm^2^); as such we denote these as B-granules (Brain-granules). As with Importin β-1 granules in motor neurons, Importin β-1 B-granules in hippocampal and cortical neurons were also evident in axons (**Figure 4C**). Quantification revealed similar abundances of cytoplasmic Importin β-1 B-granules in *C9orf72*^*+/-*^ and *C9orf72*^*-/-*^ mice in each region examined (dentate gyrus, CA3, CA1, cortex) at both 2 and 6 months of age (**Figures 4D and 4E, Table 2**). To determine whether Importin β-1 B-granules were formed during development, we performed immunofluorescence labelling of the brain from *C9orf72*^*+/-*^ or *C9orf72*^*-/-*^ neonatal mice at postnatal day 1, however no B-granules were detected (**Figure S4C**). Nor were B-granules evident in primary cortical neuron cultures from *C9orf72*^*-/-*^ mice at 7DIV (**Figure S4D**).

**Table 2.** Quantification of the abundance of Importin β-1 B-granules in *C9orf72* knockout brain

| **Average Importin β-1 B-granules per $\mu\text{m}^3$ at 2 months of age ($\times 10^{-5}$) ±SEM** |  |  |
| --- | --- | --- |
| *Region* | *C9orf72*<sup>+/-</sup> | *C9orf72*<sup>-/-</sup> |
| Dentate Gyrus | 3.43±0.88 | 4.17±1.93 |
| CA3 | 5.54±1.46 | 11.7±7.54 |
| CA1 | 25.5±3.73 | 28.1±7.92 |
| Cortex | 4.13±0.83 | 5.43±0.52 |
| Cerebellum | 0 | 0 |
| **Average Importin β-1 B-granules per $\mu\text{m}^3$ at 6 months of age ($\times 10^{-5}$)** |  |  |
| *Region* | *C9orf72*<sup>+/-</sup> | *C9orf72*<sup>-/-</sup> |
| Dentate Gyrus | 6.93±0.31 | 12.4±1.7 |
| CA3 | 4.24±2.09 | 12.9±1.1 |
| CA1 | 26.1±7.32 | 30.6±6.35 |
| Cortex | 5.11±0.72 | 4.99±0.90 |
| Cerebellum | 0 | 0 |

**Figure 4:**
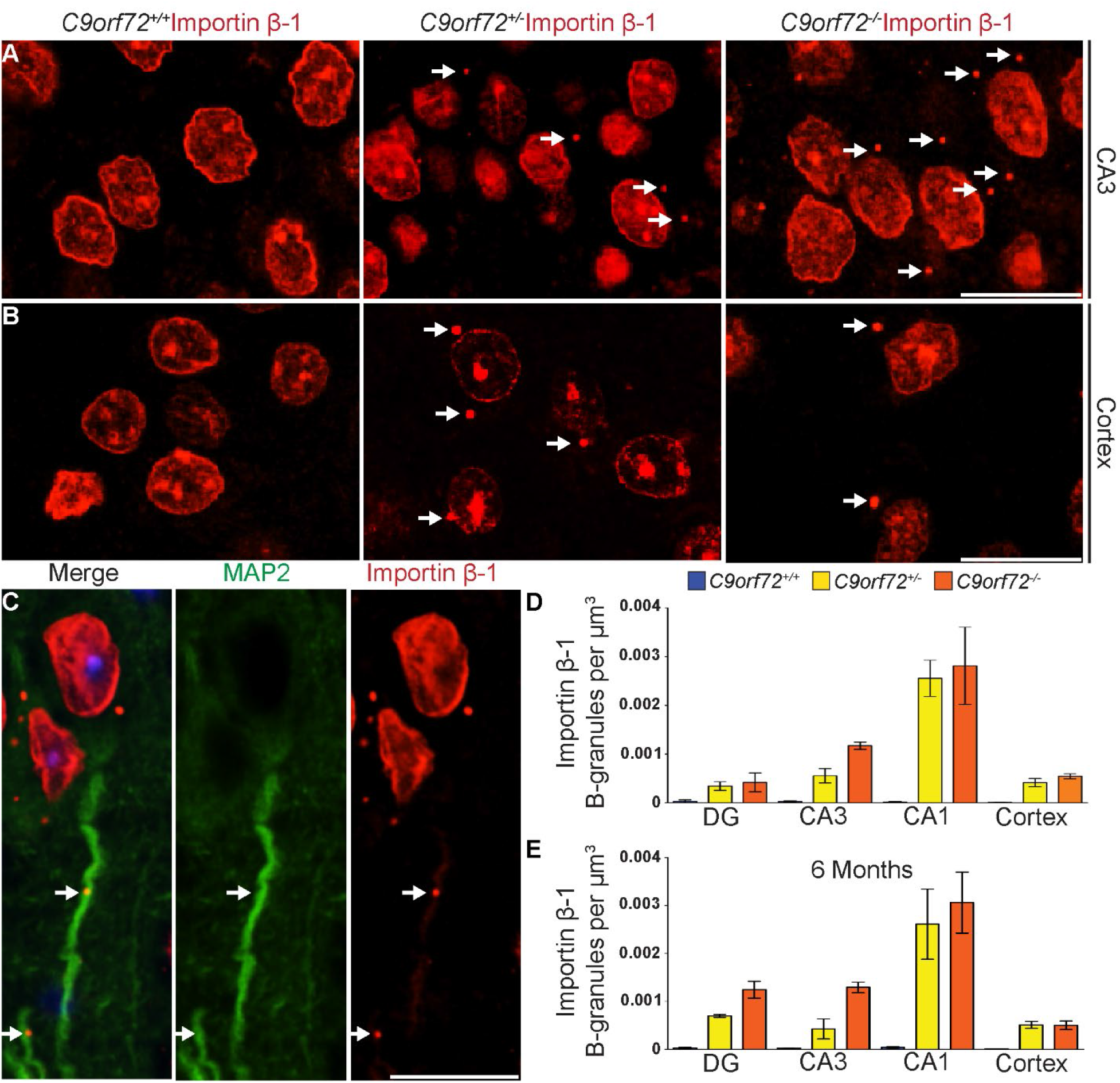
Importin β-1 granules in C9orf72 knockout mouse brain. **(A and B)** Importin β-1 immunofluorescence labeling of CA3 hippocampal region **(A)** and cortex **(B)** of *C9orf72*^*+/+*^, *C9orf72*^*+/-*^ and *C9orf72*^*-/-*^ mice at 2 months of age. White arrows denote cytoplasmic Importin β-1 granules (B-granules) present in *C9orf72* knockout neurons but absent in C9orf72^*+/+*^ neurons. **(C)** Double immunofluorescence labeling using MAP2 (green) and Importin β-1 (red) on *C9orf72*^*-/-*^ mouse brain demonstrates that some Importin β-1 B-granules localize to axons. **(D and E)** Quantification of the number of cytoplasmic Importin β-1 B-granules, normalized to the area of the region examined, in *C9orf72*^*+/+*^, *C9orf72*^*+/-*^ and *C9orf72*^*-/-*^ mice at 2 months **(D)** and 6 months of age **(E)**. n=3-4 mice per genotype. Per mouse one 63x image from DG (dentate gyrus), CA3 and CA1 was examined, and two images from cortex. Data are mean±SEM. One-way ANOVA with Bonferroni post-hoc testing did not reveal any significant differences between *C9orf72*^*+/-*^ and *C9orf72*^*-/-*^ mice at 2 or 6 months of age, however there was a trend towards significance in the DG and CA3. Scale bars = 20µm.

Whereas there were multiple Importin β-1 S- and L-granules observed in *C9orf72*^*+/-*^ and *C9orf72*^*-/-*^ motor neurons, only 1-2 Importin β-1 B-granules were observed when present in cortical and hippocampal neurons, suggesting that these granules have neuronal subtype-specific properties. Supporting this, Importin β-1 B-granules in *C9orf72*^*+/-*^ and *C9orf72*^*-/-*^ cortical and hippocampal (data not shown) neurons co-labeled with RanGAP (**Figures 5A-C**), but not with mAb414, Ubc9 or RanBP2 (**Figures S4E-G**) and there was no apparent differences in the subcellular localizations of POM121, Lamin B, Nup205, Ataxin-2 and TDP-43 in *C9orf72*^*+/+*^ or *C9orf72*^*-/-*^ hippocampal (data not shown) or cortical neurons (**Figures S4H-L**). Quantification revealed near complete colocalization of RanGAP with Importin β-1 B-granules (**Figure S5A, Table 3**), with the same regional and cell type distributions, and absence from neonates and primary cortical neurons (**Figures S5B-E**).

**Table 3.**
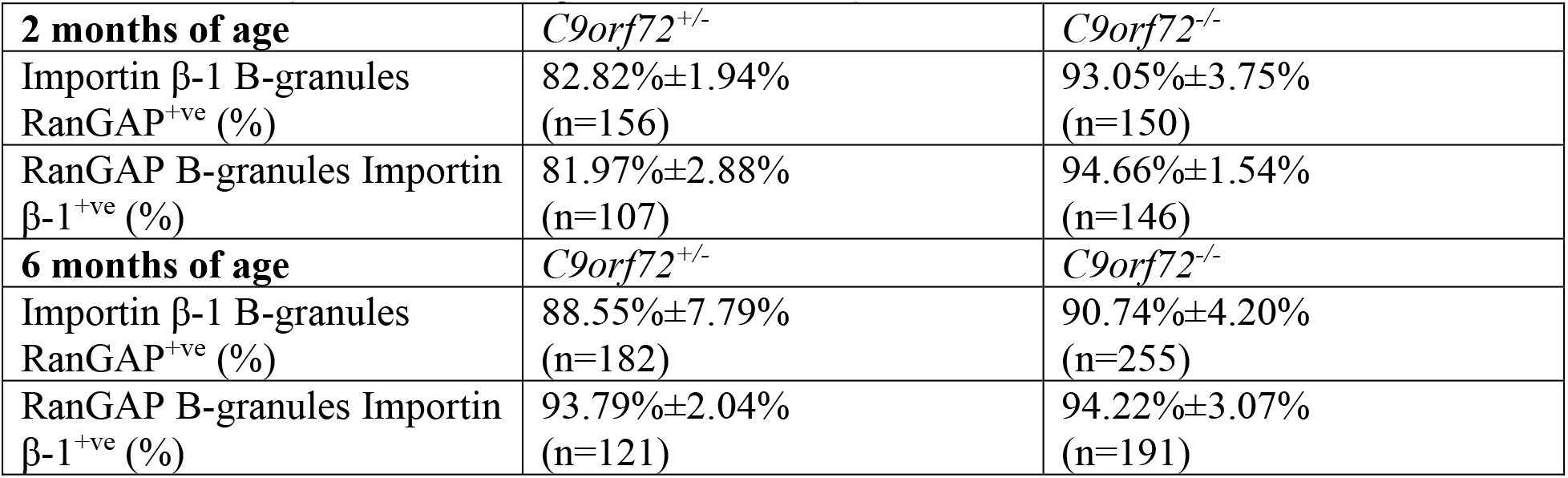
Quantification of the colocalization of Importin β-1 and RanGAP B-granules in *C9orf72* knockout cortex. (n=number of B-granules examined)

**Figure 5:**
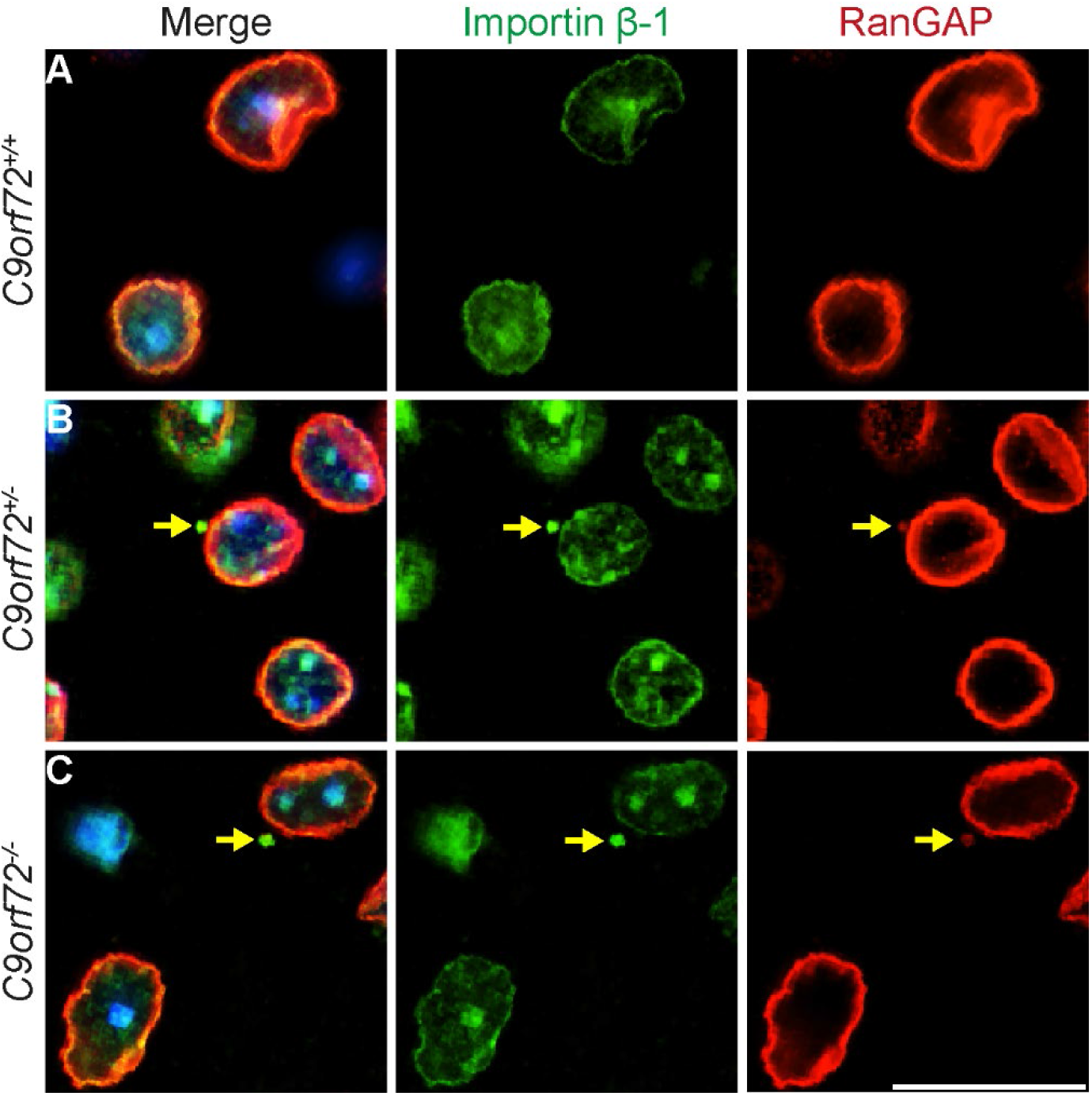
RanGAP colocalizes with cytoplasmic Importin β-1 granules in hippocampal and cortical neurons lacking C9orf72. **(A-C)** Double immunofluorescence labelling of Importin β-1 (green) and RanGAP (red) in cortical neurons of *C9orf72*^*+/+*^ **(A)**, *C9orf72*^*+/-*^ **(B)** and *C9orf72*^*-/-*^ **(C)** mice. Note colocalization of Importin β-1 B-granules with RanGAP in *C9orf72*^*+/-*^ and *C9orf72*^*-/-*^ cortical neurons (indicated with yellow arrows). DAPI nuclear stain (blue). Scale bars = 20µm.

These data demonstrate that loss of *C9orf72 in vivo* causes the formation of cytoplasmic Importin β-1 B-granules in cortical and hippocampal neurons, but not in cerebellar neurons. These granules differ in size and are compositionally distinct from Importin β-1 S- and L-granules in *C9orf72*^*+/+*^, *C9orf72*^*+/-*^ and *C9orf72*^*-/-*^ motor neurons.

### Evidence of Importin β-1 L-granules and B-granules budding from the nuclear envelope and co-association with stress granule marker G3BP1

3D deconvolution of confocal images revealed that Importin β-1 L-granules in motor neurons (**Figure 6A**) and Importin β-1 B-granules in cortical neurons (**Figure 6B**), examined through z-stacks, could be juxtaposed or contiguous with the nuclear envelope, and in some cases, the granules appeared to be tethered to the nuclear envelope (**Figures 6A and 6B**, white arrows). This gave the impression that the granules were budding from or fusing with the nuclear envelope. Interestingly, Importin β-1 was absent from the core of L-granules and B-granules, as seen from z-stacks (**Figures 6B and 6C, Figure S6A**), suggesting the presence of other components. Although appearing as predominantly spherical in structure, some Importin β-1 L-granules and B-granules were bipolar or showed protuberances (**Figure 6D, Figure S6A**), giving the impression that granules may be merging or dividing. Recent work has demonstrated that many ALS-associated proteins, including stress granule (SG) proteins and FG-Nups behave like liquid droplets^17,86–88^, of which phase separation and merging or dividing are typical properties. To investigate this possibility, we co-labelled Importin β-1 L-granules and B-granules with antibody to obligate SG protein, G3BP1. There was no co-labeling of Importin β-1 S- or L-granules with G3BP1 in *C9orf72*^*+/+*^ motor neurons (**Figure 7A**). In *C9orf72*^*+/-*^ and *C9orf72*^*-/-*^ motor neurons, G3BP1 co-localized with Importin β-1 L-granules, but not S-granules **(Figure 7B and 7C)**. Moreover, G3BP1 co-labeled Importin β-1 L-granules that appeared to be merging/dividing **(Figure 7B, Figure S6B)**, as well as those at the nuclear envelope **(Figure 7D)**. G3BP1 also colocalized with Importin β-1 B-granules in the cytoplasm **(Figure 7E)** and at the nuclear envelope **(Figure 7F)** in cortical neurons. There was near 100% colocalization of G3BP1 with Importin β-1 L-and B-granules at the nuclear envelope in both motor neurons (**Figure 7D**) and cortical neurons (**Figure 7F**), respectively. However, the frequency of G3BP1 labeling of Importin β-1 L- and B-granules in the cytoplasm, clear of the nuclear envelope, dropped to 40-60% in *C9orf72*^*+/-*^ and *C9orf72*^*-/-*^ motor neurons (**Figure 7G, Table 4**) and 20-30% in cortical neurons, respectively (**Figure 7H, Table 4**). In contrast to G3BP1, SG protein Caprin1 did not co-localize with Importin β-1 granules in *C9orf72*^*+/-*^ and *C9orf72*^*-/-*^ motor neurons or cortical neurons (**Figure S6C**), suggesting that Importin β-1 granules co-labeling with G3BP1 are not SGs.

**Table 4.** Colocalization of Importin β-1 L-granules and Importin β-1 B-granules with G3BP1 in *C9orf72* knockout mice at 2 and 6 months of age. (n=number of granules examined)

| **2 months of age** | *C9orf72*<sup>+/-</sup> | *C9orf72*<sup>-/-</sup> |
| --- | --- | --- |
| Importin β-1 L-granules<br>G3BP1<sup>+ve</sup> (%) | 64.61%±11.90%<br>(n=367) | 36.96%±24.91%<br>(n=259) |
| Importin β-1 B-granules<br>G3BP1<sup>+ve</sup> (%) | 19.44%±3.06%<br>(n=273) | 16.27%±3.33%<br>(n=344) |
| **6 months of age** | *C9orf72*<sup>+/-</sup> | *C9orf72*<sup>-/-</sup> |
| Importin β-1 L-granules<br>G3BP1<sup>+ve</sup> (%) | 45.17%±6.47%<br>(n=487) | 58.86%±6.12%<br>(n=341) |
| Importin β-1 B-granules<br>G3BP1<sup>+ve</sup> (%) | 27.49%±3.01%<br>(n=431) | 22.96%±2.94%<br>(n=384) |

**Figure 6:**
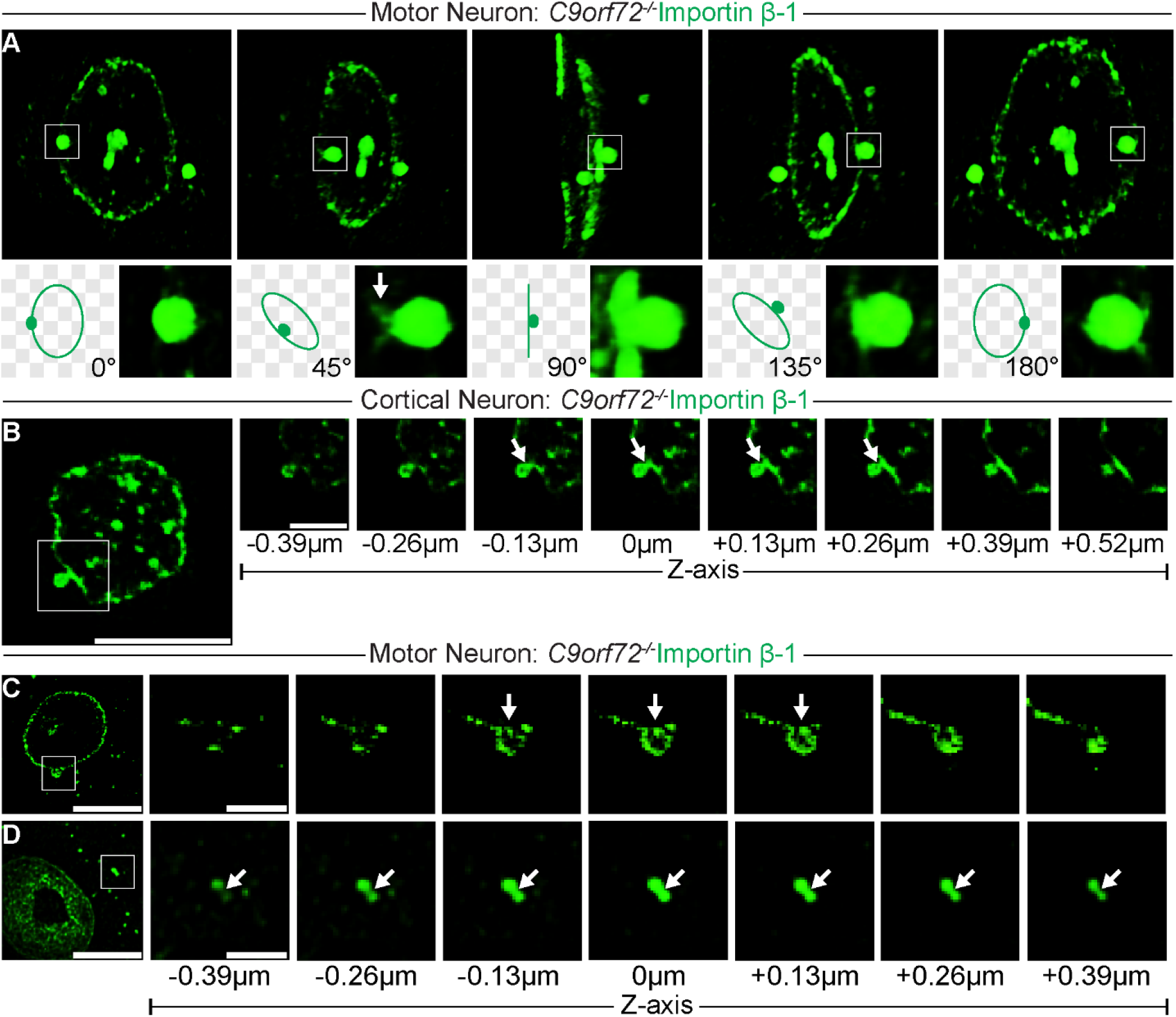
Importin β-1 granules in C9orf72 knockout neurons bud from the nuclear envelope. **(A)** Immunofluorescence labeling *C9orf72*^*-/-*^ mouse spinal cord motor neuron with Importin β-1 antibody. Confocal imaging with deconvolution and 3D rendering reveals Importin β-1 L-granule in contact with the nuclear envelope (box). Rotation (45°, 90°, 135°, 180°) shows budding of L-granules from the nuclear envelope with appearance of stalks (45°, white arrow). **(B)** Immunofluorescence labeling with Importin β-1 demonstrated that Importin β-1 B-granules in *C9orf72*^*-/-*^ cortical neurons were also found to bud from the nuclear envelope, with incremental z-stacking showing appearance of tethering (white arrows). **(C and D)** Z-stacks of deconvolved images of immunofluorescence labeling of motor neurons in *C9orf72*^*-/-*^ mouse spinal cord with Importin β-1 reveals that Importin β-1 is absent from the core of L-granules in motor neurons (**C**, white arrows) whereas some Importin β-1 L-granules are bipolar and appear as merging/dividing (**D**, white arrows). Scale bars = 10µm, 1.5µm in panel.

**Figure 7:**
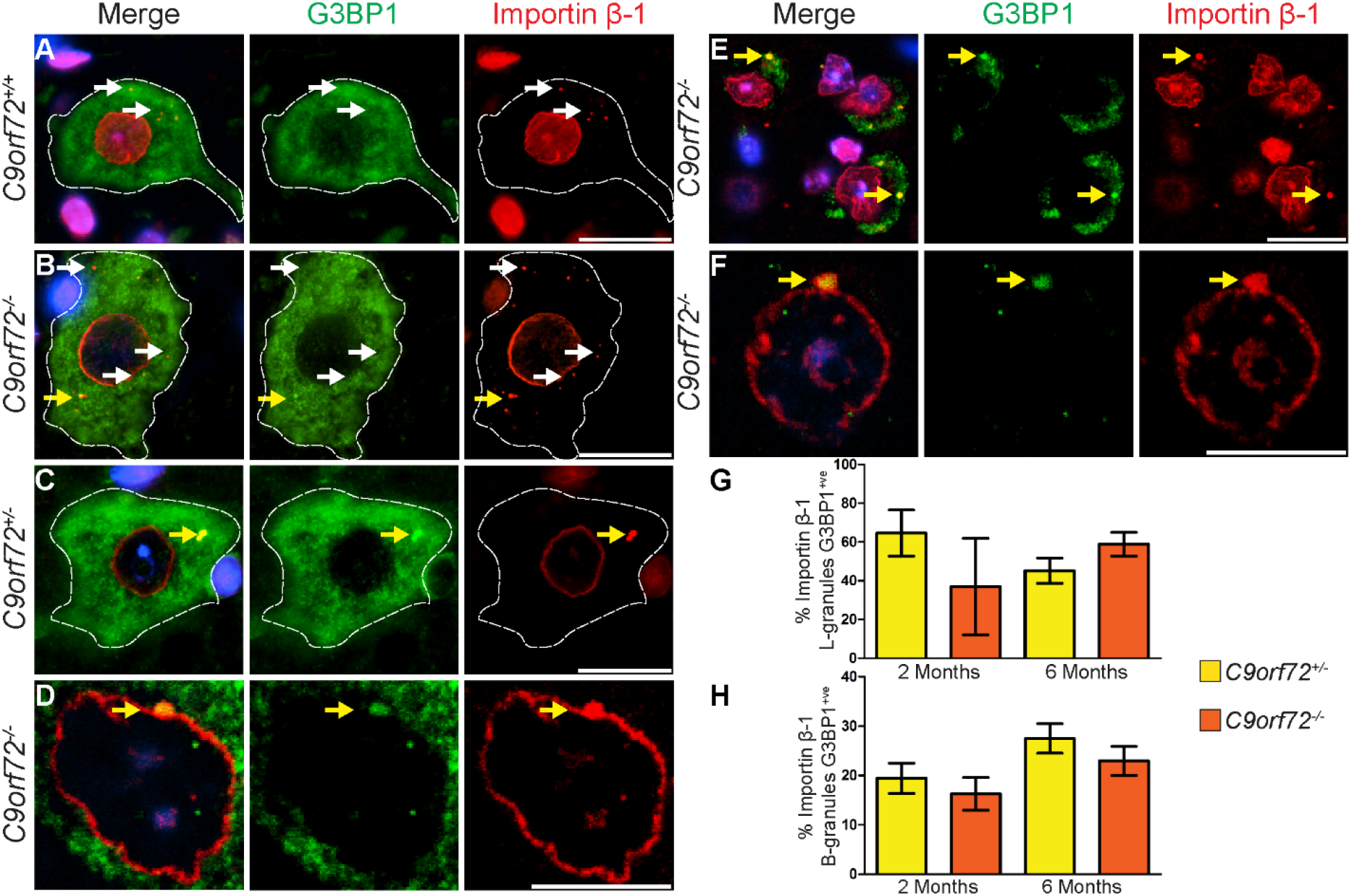
Importin β-1 granules in C9orf72 knockout motor neurons and cortical neurons colocalize with G3BP1. **(A-F**) Double immunofluorescence labeling of mouse spinal motor neurons and cortical neurons with G3BP1 (green), Importin β-1 (red) and DAPI nuclear stain (blue). Importin β-1 S-granules do not colocalize with G3BP1 in any genotype (**A and B**, white arrows) whereas Importin β-1 L-granules in *C9orf72* knockout motor neurons colocalize with G3BP1 (**B and C**, yellow arrows), including those which have the appearance of merging/dividing **(C)** and budding from the nuclear envelope **(D)**. Importin β-1 B-granules in *C9orf72*^*-/-*^ cortical neurons also displayed G3BP1 colocalization **(E)**, including during budding from the nuclear envelope **(F). (G and H)** Quantification of the percentage of G3BP1 colocalization to cytoplasmic Importin β-1 L-granules in motor neurons **(G)** and Importin β-1 B-granules in cortical neurons **(F)** of *C9orf72*^*+/-*^ and *C9orf72*^*-/-*^ mice at 2 and 6 months of age. Per genotype a minimum of 259 Importin β-1 L-granules and 273 Importin β-1 RB-granules were counted from at least n=3 mice per genotype. Data are mean±SEM. One-way ANOVA with Bonferroni post-hoc testing did not reveal any significant differences between genotypes or ages. Scale bars A, B, C, E = 20µm. Scale bars D, F = 10µm.

### Cytoplasmic Importin β-1 granules in motor and cortical neurons of *C9orf72* knockout mice colocalize with ubiquitin but not with autophagy proteins

Abnormalities of protein homeostasis, largely centered around the ubiquitin-proteasome system and autophagy, have been widely implicated in ALS pathogenesis^89^ and C9orf72 has been proposed to have roles in autophagy^47,55^. However, Importin β-1 granules (S, L and B) were not co-labeled with autophagy proteins p62 or LC3 in any genotype (**Figures S7A-D**). In contrast, approximately 5% of Importin β-1 S-granules were labeled with ubiquitin in *C9orf72*^*+/+*^ motor neurons, with no labeling of L-granules (**Figures 8A and 8D, Table 5**). Remarkably, near 100% of Importin β-1 S- and L-granules in motor neurons (**Figures 8B-D**), and 100% of Importin β-1 B-granules in cortical neurons (**Figures 8E-H**), were co-labeled with ubiquitin. The amino acid residue at which polyubiquitin chains are linked determines the fate of a ubiquitinated protein. We therefore co-labeled Importin β-1 with antibodies that recognize poly ubiquitin chains linked at K48 (which targets proteins for proteasomal degradation^90,91^) and K63 (which typically directs the protein for proteasome-independent fates or functions, including the endosomal-lysosomal pathway^90,92–96^). There was no labelling of S-granules, L-granules or B-granules with Ubiquitin-K48 in any genotype (**Figures 8I and 8J, Figures S7E and S7F**). In contrast, both Importin β-1 S- and L-granules in motor neurons and Importin β-1 B-granules in cortical neurons of *C9orf72*^*+/-*^ and *C9orf72*^*-/-*^ mice, were labeled with Ubiquitin-K63 (**Figures 8K and 8L**), including those at the nuclear envelope and those that appeared to be merging/dividing (**Figures S7I and S7J**).

**Table 5.** Quantification of the colocalization of ubiquitin to Importin β-1 S-granules, L-granules and B-granules in *C9orf72* knockout mice at 6 months of age. NA=not applicable. (n=number of granules examined)

| **Importin β-1 granules** | *C9orf72*<sup>+/+</sup> | *C9orf72*<sup>+/-</sup> | *C9orf72*<sup>-/-</sup> |
| --- | --- | --- | --- |
| Importin β-1 S-granules ubiquitin<sup>+ve</sup> (%) | 5.37%±0.93%<br>(n=246) | 82.64%±5.10%<br>(n=208) | 92.33%±3.56%<br>(n=351) |
| Importin β-1 L-granules ubiquitin<sup>+ve</sup> (%) | 0%±0%<br>(n=8) | 100%±0%<br>(n=80) | 97.62%±2.38%<br>(n=58) |
| Importin β-1 B-granules ubiquitin<sup>+ve</sup> (%) | NA | 100%±0%<br>(n=76) | 100%±0%<br>(n=111) |

**Figure 8:**
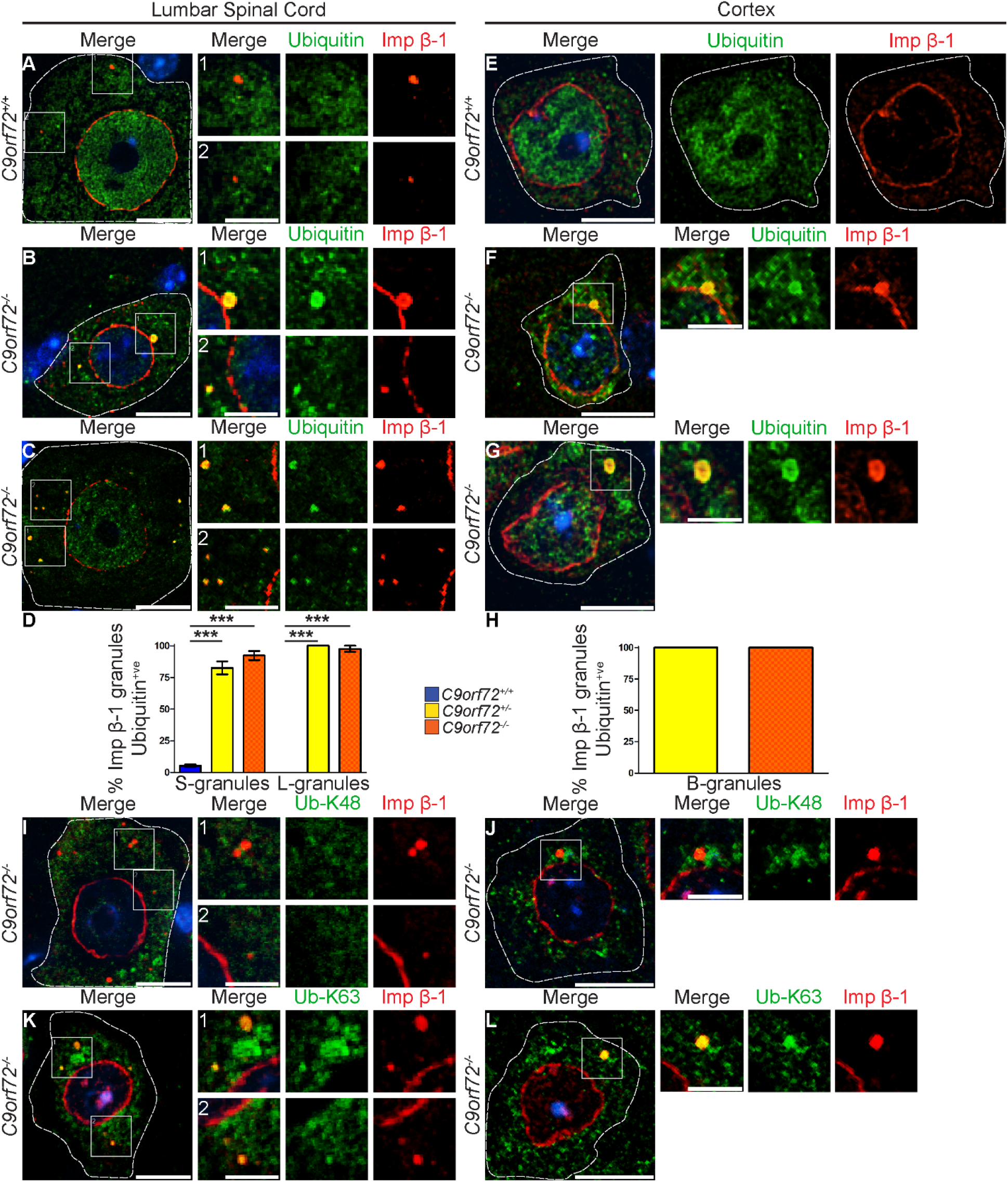
Ubiquitin colocalizes to cytoplasmic Importin β-1 granules in motor neurons and cortical neurons of C9orf72 knockout mice. **(A-H)** Double immunofluorescence labeling, followed by colocalization quantification, for ubiquitin (green), Importin β-1 (red) and DAPI nuclear stain (blue) in mouse spinal motor neurons (**A-D**) and cortical neurons **(E-H)**. *C9orf72*^*+/+*^ motor neurons demonstrated a lack of colocalization of ubiquitin to Importin β-1 L-granules (panel 1) or S-granules (panel 2) at 6 months of age **(A)**, whereas *C9orf72*^*-/-*^ motor neurons demonstrated ubiquitin colocalization to both Importin β-1 L-granules, during budding (**B**, panel 1) and in the cytoplasm (**C**, panel 1), and Importin β-1 S-granules (**B**, panel 2; **C**, panel 2). Of note, ubiquitin colocalization does not occur with typical Importin β-1 staining at the nuclear envelope. **(D)** Quantification of ubiquitin^+ve^ Importin β-1 S-granules and L-granules in *C9orf72*^*+/+*^, *C9orf72*^*+/-*^ and *C9orf72*^*-/-*^ motor neurons at 6 months of age demonstrated little or no colocalization to *C9orf72*^*+/+*^ Importin β-1 S-granules or L-granules, and extensive colocalization with S-granules and L-granules in *C9orf72*^*+/-*^ and *C9orf72*^*-/-*^ motor neurons. Importin β-1 and ubiquitin colabeling in *C9orf72*^*+/+*^ cortical neurons demonstrated expected subcellular localizations **(E)**, whereas Importin β-1 B-granules in *C9orf72*^*-/-*^ cortical neurons displayed ubiquitin colocalization during budding **(F)** and in the cytoplasm **(G). (H)** Quantification of ubiquitin^+ve^ Importin β-1 RB-granules in *C9orf72*^*+/-*^ and *C9orf72*^*-/-*^ cortical neurons demonstrated complete colocalization to every B-granule examined. **(I-L)** Double immunofluorescence labelling of *C9orf72*^*-/-*^ tissue for ubiquitin chains linked at the K48 residue (Ub-K48, green) or K63 residue (Ub-K63, green) and Importin β-1 (red) demonstrates a lack of colocalization between any Importin β-1 granules and Ub-K48 in motor neurons **(I)** and cortical neurons **(J)**, but clear colocalization of Ub-K63 with both Importin β-1 L-granules (**K**, panel 1) and S-granules (**K**, panels 1 and 2) in motor neurons and Importin β-1 B-granules in cortical neurons **(L)**. For quantification, n=3 mice per genotype, with n=208-351 S-granules and n=8-80 L-granules examined per genotype, and 3-5 63x images of cortex were scored per animal examining n=76-101 B-granules. Data are mean±SEM. Neuronal cell bodies are denoted by white dotted line. One-way ANOVA with Bonferroni post-hoc testing was used for statistical analysis of ubiquitin^+ve^ granules in lumbar spinal cord motor neurons. p<0.001=***. Scale bars = 10µm, 5µm inset. Imp β-1 = Importin β-1.

## Discussion

Here we have identified a neuronal phenotype in *C9orf72*^*+/-*^ and *C9orf72*^*-/-*^ mice. We have shown that loss of C9orf72 disrupts the neuronal Ran-GTPase gradient, both *in vitro* and *in vivo*, and that this was associated with defects in NCT. We identified two types of cytoplasmic Importin β-1 granules (S-granules and L-granules) that co-localized with NPC proteins in motor neurons of *C9orf72*^*+/+*^ mice. In *C9orf72*^*+/-*^ and *C9orf72*^*-/-*^ motor neurons, the abundance of Importin β-1 S- and L-granules increased and there was a loss of co-labeling with NPC proteins. Importin β-1 granules were absent in *C9orf72*^*+/+*^ mouse brain, but Importin β-1 B-granules, compositionally distinct from those observed in motor neurons, were present in cortical and hippocampal neurons of *C9orf72*^*+/-*^ and *C9orf72*^*-/-*^ knockout mice and co-labeled with RanGAP. Importin β-1 L- and B-granules appeared to be generated through budding from the nuclear envelope and were co-labeled with G3BP1 and K63-linked ubiquitin. These findings link *C9orf72* haploinsufficiency to similar defects in NCT and NPC proteins observed for RNA and DPR toxicities arising from the *C9orf72* repeat expansions.

Our previous work suggested that C9orf72 could potentially function in NCT through interactions with Ran-GTPase and Importin β-1^29^. This is supported by other studies showing that C9orf72 and its binding partner SMCR8 can interact with Importin β-1 and regulators of the Ran-GTPase cycle^46,65,66^. Here we have demonstrated that loss of C9orf72 causes an increase in cytoplasmic Ran-GTPase in HeLa cells, primary motor and cortical neurons, and in spinal motor neurons *in vivo*. This disruption of the Ran-GTPase gradient was associated with defects in NCT. Prior studies have shown that impairments of the Ran-GTPase gradient can be caused by a variety of stressors^73,97,98^ and has been demonstrated in numerous *in vitro* and *in vivo* ALS/FTD models^12,17,24,36,73,97^. Associated with disruption of the Ran-GTPase gradient, we identified different types of cytoplasmic Importin β-1 granules *in vivo* that exhibited genotype and neuronal-subtype specific properties.

In motor neurons of *C9orf72*^*+/+*^ mice we observed two populations of cytoplasmic Importin β-1 granules that could be differentiated based on size: S-granules (∼0.15µm^2^), which were abundant, and L-granules (∼0.30µm^2^), which were rare. Both types of granules were co-labeled with antibodies to FG-nucleoporins (mAb414), RanGAP, Ubc9 and RanBP2 (the RanBP2 complex), but not with POM121, suggesting that these structures may be related to annulate lamellae pore complexes (ALPCs)^83–85,99–101^. ALPCs are typically observed in rapidly dividing cells, are morphologically similar to NPCs, and are localized to specialized regions of the endoplasmic reticulum^84,102^. Despite being described over 60 years ago^100^, there is no agreed consensus of their function. ALPCs have been proposed to regulate NCT and NPC assembly^99^, act as intermediate docking sites for nuclear import complexes^99^, sites for nuclear export complex dissociation^99^, or act as a stockpile of NPCs for daughter cells during embryogenesis^103– 106^. Strikingly, in *C9orf72* knockout motor neurons there was a significant increase in the total number of cytoplasmic Importin β-1 granules, however there was a loss of co-labeling with mAb414, RanGAP, Ubc9 and RanBP2. Studies in *Drosophila* oocytes, have demonstrated that early steps in ALPC biogenesis involve phase separation of FG-Nups, including RanBP2, to generate compositionally diverse biomolecular condensates (granules) that vary in size, and merge/divide to exchange contents. Importin β-1 binding to FG-Nups in these granules, promoted by RanGDP, prevents their assembly into ALPCs, whereas elevated RanGTP dissociates Importin β-1, allowing the FG-Nups to assemble^104,107,108^. The Importin β-1 granules we have observed appeared to have properties of merging/dividing, consistent with the granules observed in *Drosophila* oocytes. Furthermore, it is possible that the loss of co-labelling of Importin β-1 granules in *C9orf72* knockout motor neurons with mAb414, RanGAP, Ubc9 and RanBP2 is due to elevated levels of RanGTP. However, converse to the study in *Drosophila* oocytes, we did not observe an increase in ALPCs, but rather an increase in Importin β-1 granules. As such the identity and mechanism of genesis of Importin β-1 granules in *C9orf72*^*+/-*^ and *C9orf72*^*-/-*^ motor neurons remains to be elucidated.

In contrast to motor neurons, cytoplasmic Importin β-1 granules were not evident in cortical or hippocampal neurons of *C9orf72*^*+/+*^ mice. Cytoplasmic Importin β-1 granules were however observed in cortical and hippocampal neurons of both *C9orf72*^*+/-*^ and *C9orf72*^*-/-*^ mice, but not in cerebellar neurons. These granules were significantly larger than those observed in motor neurons (0.44µm^2^±0.02µm^2^, versus L-granules: 0.30µm^2^±0.02µm^2^) and were less abundant (1-2 per cortical/hippocampal neuron versus up to 40 in motor neurons). Interestingly, B-granules were co-labeled with RanGAP, but not with mAb414, Ubc9 or RanBP2, indicating that these granules are compositionally different from those observed in *C9orf72* knockout mouse motor neurons. The compositional diversity of Importin β-1granules suggests that loss of C9orf72 has differential effects on vulnerable neuronal subpopulations, which may be a determinant of whether *C9orf72* mutation carriers develop ALS, FTD or ALS/FTD.

A commonality between L-granules in motor neurons and B-granules in cortical neurons was that each could be observed juxtaposed to the nuclear envelope, with the appearance of budding, and granules tethered to the envelope. Protuberances from the nuclear envelope (referred to in the literature as herniations, blebs and/or buds^109,110^) have been proposed to be involved with, in addition to other roles, NPC assembly and NPC quality control/surveillance^109^. Nuclear budding could also have deleterious consequences, for example inducing toxicity through DNA damage and/or abnormal dispersal of nuclear proteins into the cytoplasm^111–114^. It is interesting that despite the differences in composition, both L- and B-granules in *C9orf72* knockout neurons were co-labeled with G3BP1, and all granules (including S-granules) were co-labelled with K63-linked ubiquitin. This is, to our knowledge, the first time that such types of nuclear buds have been described, and their appearance is related to loss of C9orf72.

There was near complete overlap of G3BP1 with L- and B-granules at the nuclear envelope, with reduced co-labeling in the cytosol, and no co-labeling observed for S-granules. Despite G3BP1 colocalization, there was no co-labeling with Caprin1, suggesting that Importin β-1 granules are not stress granules. Nevertheless, aspects of stress granule biology may underlie G3BP1 colocalization to Importin β-1 granules, particularly the ability of G3BP1 to phase separate^87^. Phase separation of proteins is dependent upon exceeding a concentration threshold and for stress granule formation, core constituents cooperate to set threshold concentrations^115^. FG-Nups have the ability to phase separate^86,116^, and phase separation is a notable property of the RanBP2 condensates that are precursors for ALPCs^104^. Notably, Importin β-1 acts as a chaperone and can regulate the phase-separation properties of FG-Nups^117–119^. It is notable that G3BP1 contains a highly conserved N-terminal nuclear transport factor 2 (NTF2)-like domain, which interacts with nucleoporin FxFG repeats^120^ and has been suggested to have a role in nuclear shuttling. This would be consistent with the near 100% co-localization of G3BP1 with Importin β-1 L- and B-granules at the nuclear envelope, although these granules are not co-labeled with FG-Nups. C9orf72 has proposed roles in autophagy^47,53,55^, however we did not find colocalization of Importin β-1 granules with autophagy proteins p62 or LC3. In contrast, Importin β-1 S-, L- and B-granules were co-labelled with ubiquitin in *C9orf72*^*+/-*^ and *C9orf72*^*-/-*^ motor and cortical neurons. The ubiquitin chains that colocalized to Importin β-1 granules in *C9orf72*^*+/-*^ and *C9orf72*^*-/-*^ neurons were linked by K63 residues, indicating a non-proteasomal fate^90^. Interestingly, it has recently been demonstrated that G3BP1 in stress granules caused by heat shock is K63-linked ubiquitinated^121^. K63-linked ubiquitination of G3BP1 is essential for stress granule disassembly, promoting recruitment of p97/VCP (valosin-containing protein) through interaction with FAF2 at the endoplasmic reticulum^121^. It is possible that G3BP1 is the is K63-linked ubiquitinated target in L- and B-granules, however, although there is 100% co-localization of G3BP1 and K63-linked ubiquitination at the nuclear envelope, there is reduced labeling with G3BP1 in the cytosol, and although near 100% of S-granules are K63-linked ubiquitinated, none are co-labeled with G3BP1. As such the protein target of the K63-linked ubiquitination is unknown, but similar processes involving dis-segregation of granule constituents akin to the mechanism observed for G3BP1 in heat shock induced stress granules may be involved.

C9orf72 has been linked with several cellular processes, including regulation of actin dynamics^46^, autophagy initiation^47,48,50,55^, endosomal trafficking^56,122–124^, lysosomal biogenesis^56,125^, immune regulation^60,61,63,75–77^, synaptic function^42,57,126^, and, as we have demonstrated here, nucleocytoplasmic transport. Our findings of genotype and neuronal subtype dependent differences in Importin β-1 granules between *C9orf72*^*+/+*^ and *C9orf72* knockout mice suggests that C9orf72 function is context dependent. The C9orf72-SMCR8 complex has been demonstrated to act as a GTPase-activating protein (GAP) for small GTPases Rab8a^52^, Rab11a^52^ and ARF1^51,127^. Although currently identified small GTPase targets of C9orf72 are not implicated in NCT or Importin β-1 function, our data clearly demonstrate dysfunction of these pathways in the context of C9orf72 reduction. It remains to be established if C9orf72 works directly or indirectly on these pathways. We note that current knowledge of C9orf72 interactions and functions are largely based on *in vitro* findings from non-neuronal cells^46,48,50,53,65,66,122^, and therefore may not delineate neuronal-specific properties.

In conclusion, our findings demonstrate that reduction or complete loss of C9orf72 causes changes in the Ran-GTPase gradient and Importin β-1 abnormalities in the neuronal subtypes affected in ALS and FTD *in vivo*. Due to the absence of neurodegeneration in *C9orf72* knockout mice^59^, Ran-GTPase mislocalization and Importin β-1 granules are seemingly innocuous under normal conditions. As such, we speculate that these are early events in disease pathogenesis that precede other forms of neuronal dysfunction and may act as a sensitizer to neurodegeneration or ‘first hit.’ Published evidence has demonstrated that the *C9orf72* gain-of-function mechanisms disrupt NCT, with genetic modifier screens identifying Importin β-1^40^, other karyopherins (KPNA4, KPNA4)^40^, or RanGAP^24,40^ as enhancers of DPR or G_4_C_2_ toxicity. Moreover, Importin β-1 has been shown to bind arginine-rich DPRs^35,41^ and suppress their toxic effects^41^. Therefore, we establish NCT and Importin β-1 dysfunction as a phenotype arising from loss of C9orf72 that could act as a convergence point for additive or synergistic effects in the context of *C9orf72* gain-of-function mechanisms, leading to neurodegeneration.

## Acknowledgements

We thank Don Cleveland (UCSD) and Clotilde Lagier-Tourenne (UMass) for *C9orf72* knockout mice. Thanks to Paul McKeever (University of Toronto), Christine Vande Velde (University de Montreal), Valerie Wallace (UHN, University of Toronto), and Joel Watts (University of Toronto) for critical reading of the manuscript. This work was funded by an ALS Canada-Brain Canada Hudson Grant and the James Hunter ALS Initiative. JR holds the James Hunter Family Chair in ALS Research. PM was supported by the Christopher Chiu Postdoctoral Fellowship from ALS Double Play. TMD is supported by funding of the ALS-RAP initiative (https://als-rap.org/) through ALS Canada, the ALS Alliance and the Motor Neuron Disease Association.

## Author Contributions

PM and JR planned and designed the research. PM performed all research. AL administered mouse colonies and planned experiments. ZY and TMD generated and provided the *C9orf72* knockout HeLa cells. JR acquired funding. PM and JR wrote the manuscript.

