## Supplementary Figures for "Loss of C9orf72 perturbs the Ran-GTPase gradient and generates compositionally diverse cytoplasmic Importin β-1 granules in motor and cortical neurons *in vivo*"

**Supplemental Information**

**Figure S1**

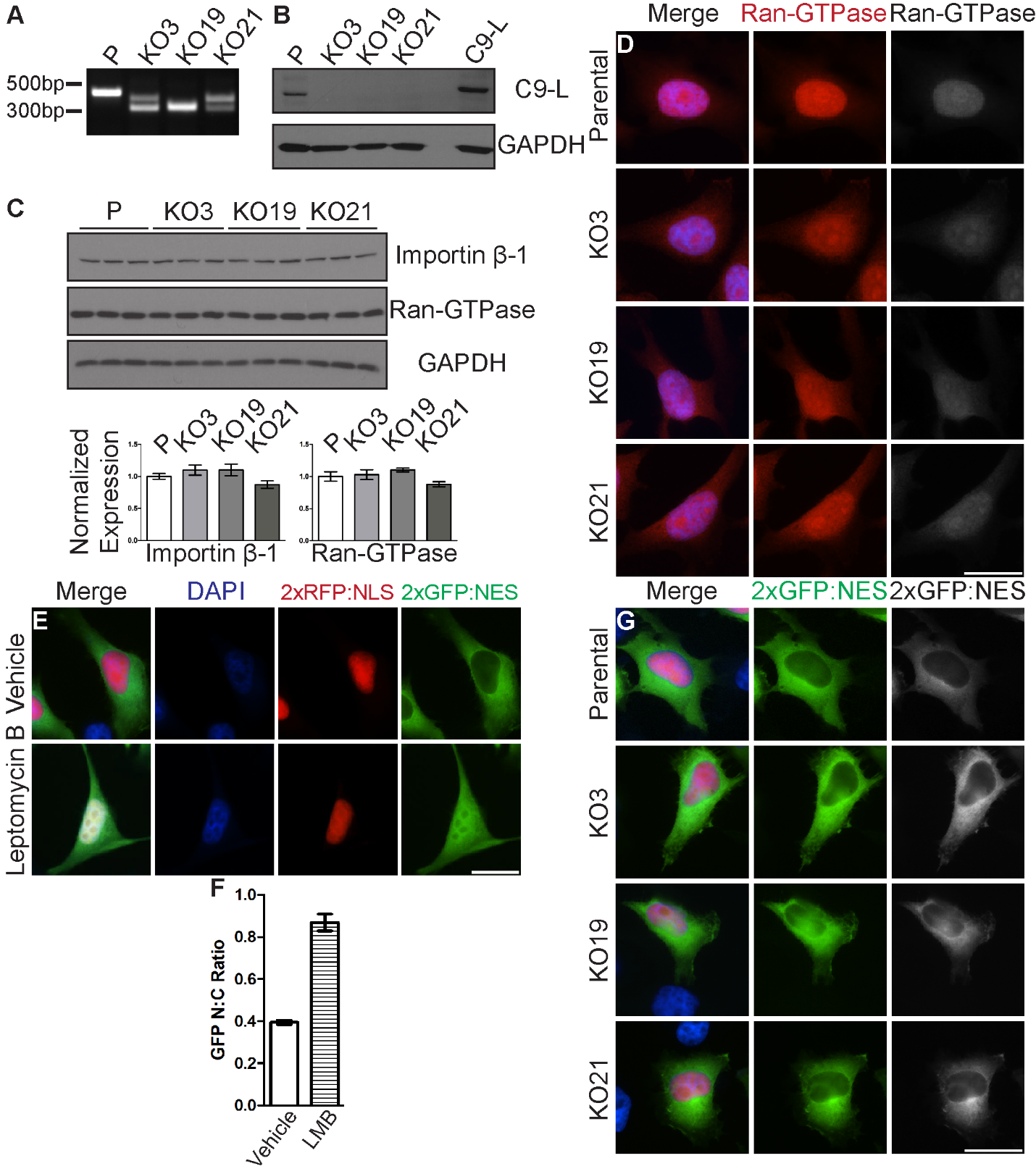

***S1: CRISPR-Cas9 mediated knockout of C9orf72 in HeLa cells***

**(A)** Polymerase chain reaction (PCR) of HeLa cell genomic DNA using primers located in the first intron of *C9orf72*. Smaller DNA fragments indicate deletions in the *C9orf72* knockout lines (KO3, KO19, KO21) compared to the parental line (P). **(B)** Western blotting of cell lysates shows the presence of C9-L in parental but not *C9orf72* knockout lines. Parental cells transfected with a construct expressing C9-L was used as positive control (lane labelled C9-L). Blots probed with C9orf72 antibody (GTX634482). **(C)** Levels of Importin β-1 and Ran-GTPase, normalized to GAPDH, were unchanged in *C9orf72* knockout versus parental (P) cells. **(D)** Immunofluorescence staining of Ran-GTPase (red) in parental and *C9orf72* knockout HeLa cells, related to Figure 1A and 1B. Note increased cytoplasmic mislocalization of Ran-GTPase in the knockout cell lines. **(E and F)** Parental cells (P) expressing nucleocytoplasmic transport reporter 2xGFP:NES-IRES-2xRFP:NLS show predominantly cytoplasmic GFP (green) and nuclear RFP (red) localization under normal conditions **(E)**. Treatment of cells with nuclear export inhibitor Leptomycin B causes accumulation of GFP in the nucleus. Quantification of GFP N:C shows an increase following Leptomycin B (LMB) treatment **(F)**. **(G)** Representative images of parental and *C9orf72* knockout HeLa cells expressing pLVX-EF1alpha-2xGFP:NES-IRES-2xRFP:NLS reporter construct, related to Figure1C and 1D. DAPI nuclear stain (blue). Scale bars = 20µm.

**Figure S2**

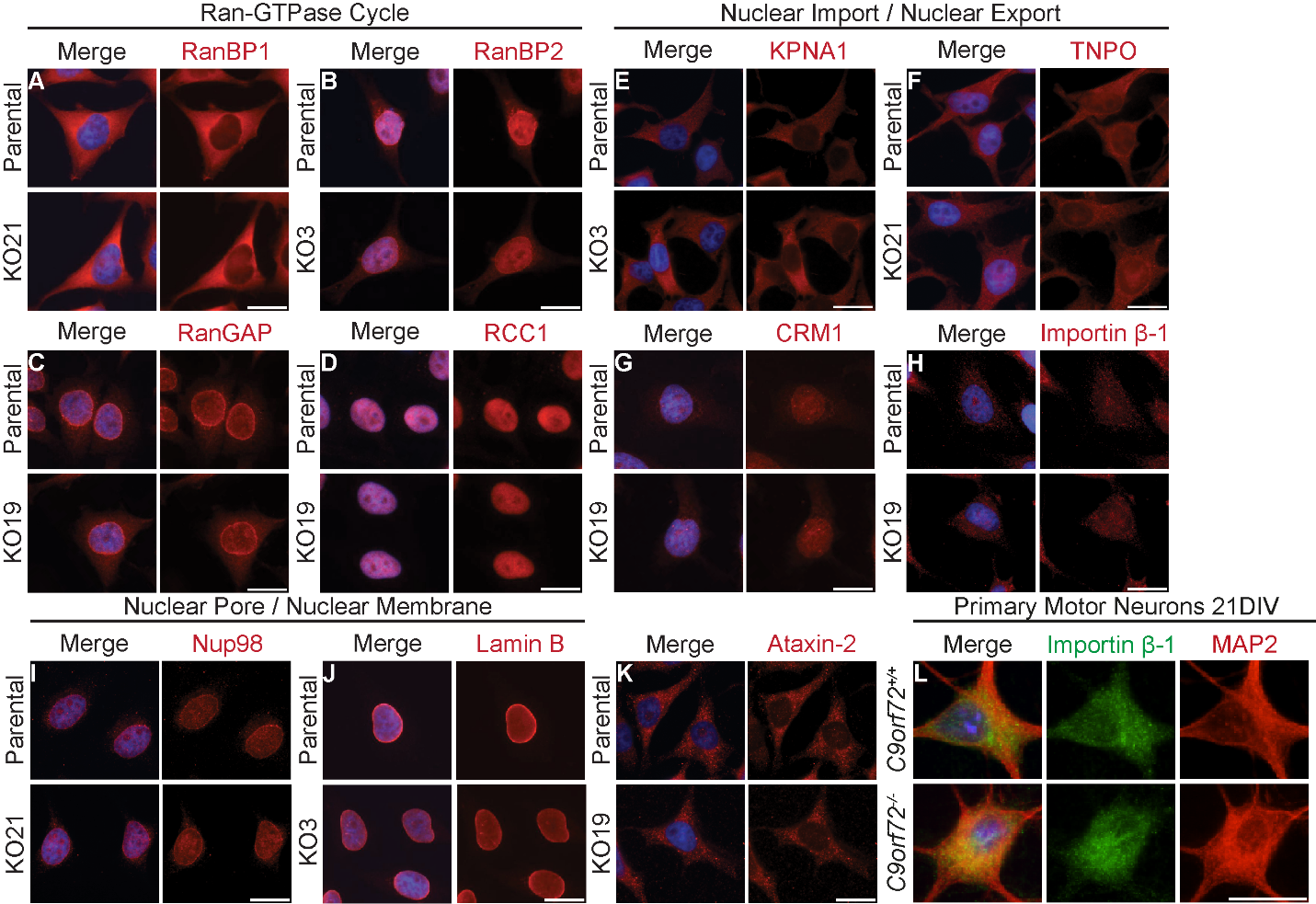

***S2: No gross mislocalization of proteins involved in NCT in C9orf72 knockout HeLa cells or primary motor neurons***

**(A-K)** Parental and *C9orf72* knockout HeLa cells immunofluorescently labeled with antibodies to proteins involved in the Ran-GTPase cycle: RanBP1 **(A)**, RanBP2 **(B)**, RanGAP **(C)**, RCC1 **(D)**; nuclear import or export: KPNA1 **(E)**, TNPO **(F)**, CRM1 **(G)**, Importin β-1 **(H)**; the nuclear membrane/nuclear pore: Nup98 **(I)** and Lamin B **(J)**, and Ataxin-2 (**K**). For each protein examined, localization was assessed in parental line and each *C9orf72* knockout line (KO3, KO19, KO21) and representative images randomly selected. **(L)** Primary motor neurons from *C9orf72^+/+^* and *C9orf72^-/-^* mice at 21 days *in vitro* (DIV) labelled with Importin β-1 (green) and neuronal marker MAP2 (red) did not show mislocalization of Importin β-1. DAPI nuclear stain (blue). Scale bars = 20µm.

**Figure S3**

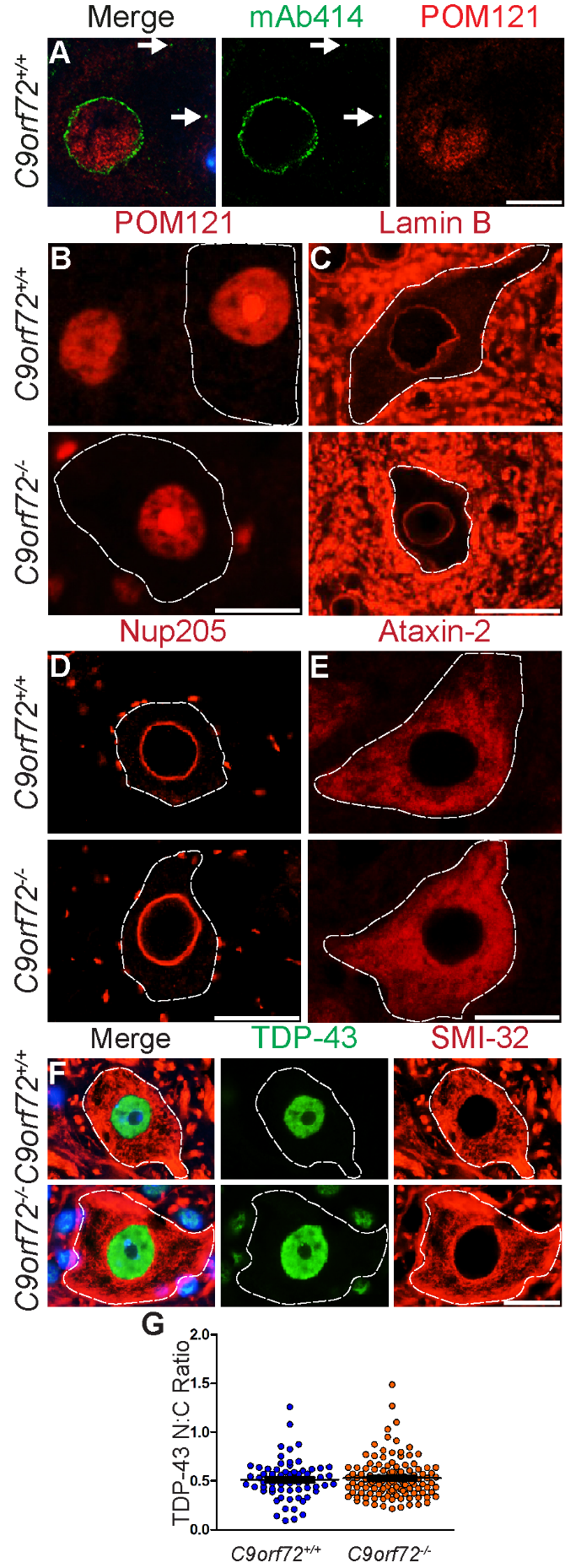

***S3: No abnormalities of selected NCT-associated proteins in C9orf72^-/-^ motor neurons***

**(A)** Double immunofluorescence labeling of mAb414 (green) and POM121 (red) in *C9orf72^+/+^* motor neurons did not display any evidence of POM121 colocalization to cytoplasmic mAb414 granules. **(B-E)** Immunofluorescence labelling of *C9orf72^+/+^* and *C9orf72^-/-^* lumbar spinal cord did not reveal any gross mislocalizations of nuclear pore protein POM121 **(B)**, nuclear lamina protein Lamin B **(C)**, nuclear pore protein Nup205 which also labelled the neuropil **(D)** or Ataxin-2 **(E)**. **(F)** Double immunofluorescence labelling of lumbar spinal cord motor neurons of *C9orf72^+/+^* and *C9orf72^-/-^* mice with antibodies to TDP-43 (green) and motor neuron marker SMI-32 (red). **(G)** Quantification of TDP-43 N:C ratio in lumbar motor neurons did not reveal any significant differences between *C9orf72^+/+^* and *C9orf72_­_^-/-^* mice at 6 months of age. Data are mean±SEM. One-way ANOVA with Bonferroni post-hoc testing was used for statistical analysis. Motor neuron cell bodies are denoted by white dotted line. DAPI nuclear stain (blue). Scale bar A = 10µm. Scale bars B-E = 20µm.

**Figure S4**

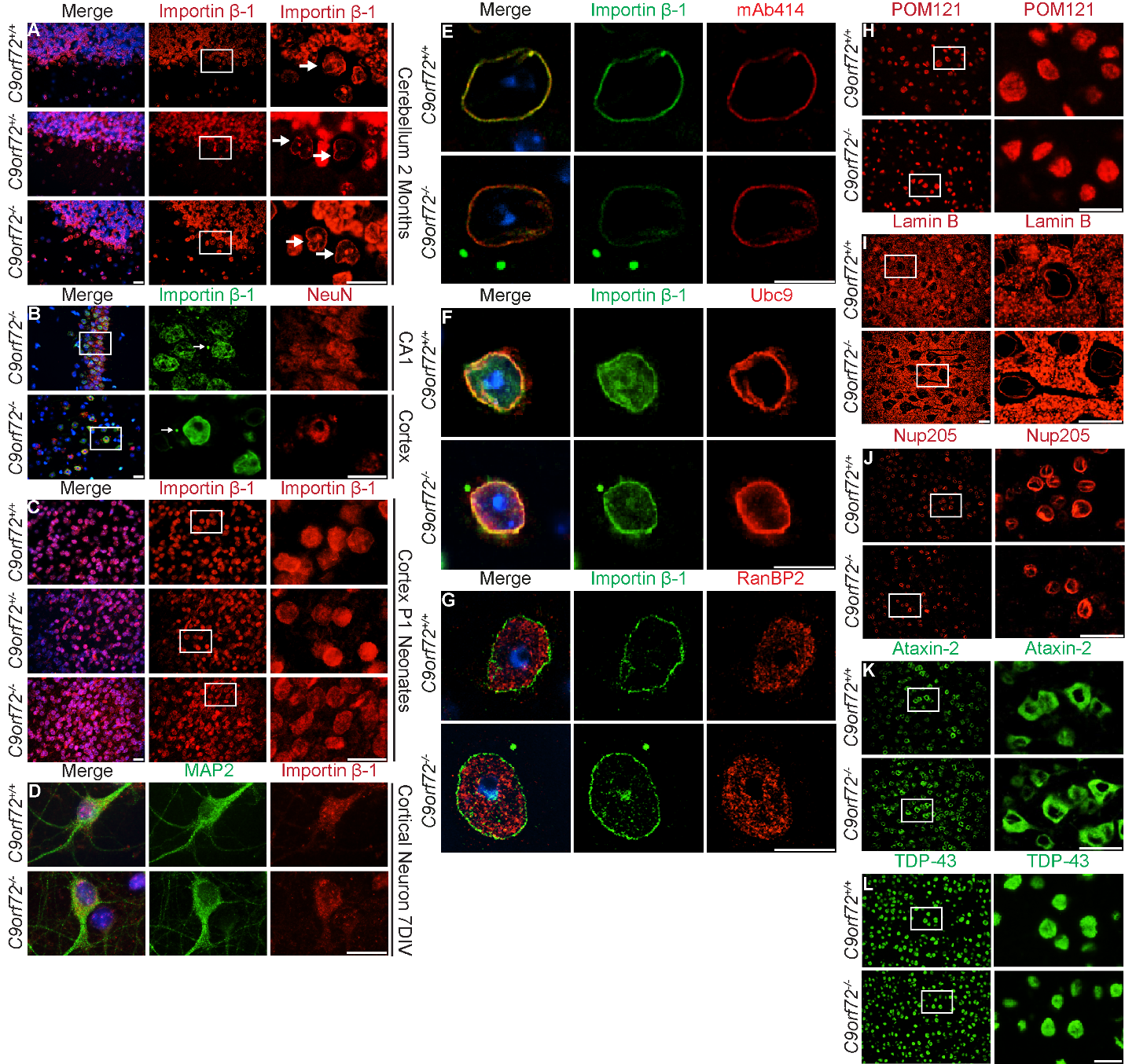

***S4: Co-labeling of Importin β-1 B-granules with NCT proteins***

**(A)** Importin β-1 immunofluorescence labeling reveals absence of Importin β-1 B-granules in Purkinje neurons (white arrows) in *C9orf72^+/+^*, *C9orf72^+/-^* and *C9orf72^-/-^* mice at 2 months of age. **(B)** Double immunofluorescence labelling of Importin β-1 (green) and NeuN (red) in CA1 and cortex of *C9orf72^-/-^* mice demonstrates the neuronal-identity of cells containing cytoplasmic Importin β-1 B-granules (white arrows). **(C)** Importin β-1 B-granules were absent in neonatal cortex (P1) of all gentotypes. **(D)** Double immunofluorescence of Importin β-1 (red) and MAP2 (green) in *C9orf72^+/+^* and *C9orf72^-/-^* primary cortical neurons at 7DIV did not reveal any Importin β-1 B-granules. **(E-G)** Double immunofluorescence labeling of Importin β-1 (green) with mAb414 **(**red, **E**), Ubc9 **(**red, **F)** and RanBP2 (red, **G**) in cortical neurons of *C9orf72^+/+^* and *C9orf72^-/-^* mice at 6 months of age, did not reveal cytoplasmic granules or colocalization with Importin β-1 B-granules in any genotype. **(H-L)** Immunofluorescence labelling of *C9orf72^+/+^* and *C9orf72^-/-^* cortical neurons did not reveal any gross mislocalizations of nuclear pore protein POM121 **(H)**, nuclear lamina protein Lamin B **(I)**, nuclear pore protein Nup205 **(J)**, Ataxin-2 **(K)** or TDP-43 **(L)**. DAPI nuclear stain (blue). Scale bars A-D, H-L = 20µm. Scale bars E-G = 10µm. DIV = days *in vitro*, P1 = postnatal day 1.

**Figure S5**

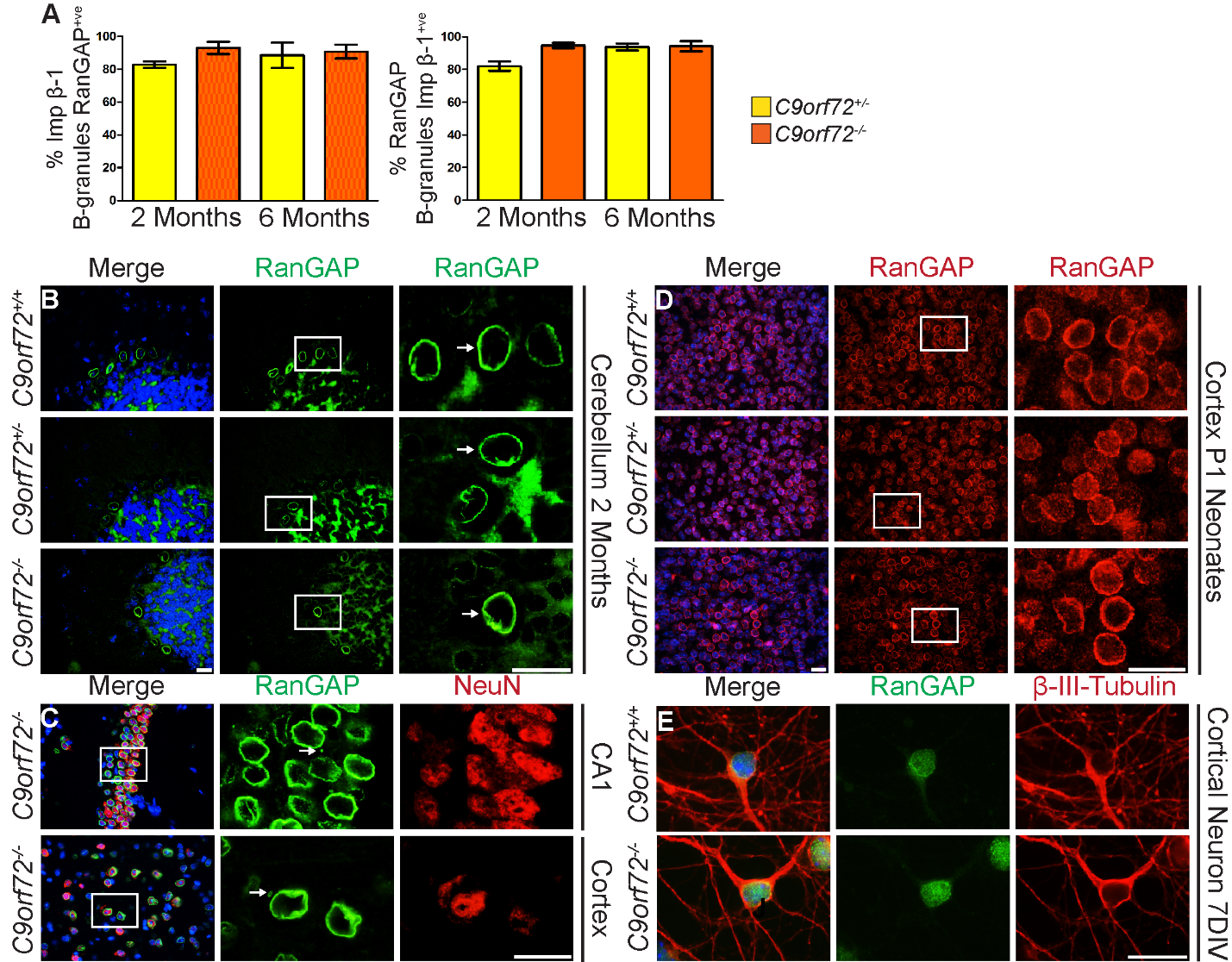

***S5: Cytoplasmic* Importin β-1 and *RanGAP show nearly complete colocalization to B-granules in cortical neurons***

**(A)** Quantification of the percentage of Importin β-1 B-granules positive for RanGAP, and percentage of RanGAP B-granules positive for Importin β-1 in the cortex of *C9orf72^+/-^* and *C9orf72^-/-^* mice at 2 and 6 months of age. n=3-4 mice per genotype. A minimum of 107 B-granules were examined per labeling, per genotype. **(B-E)** Immunofluorescence demonstrates that RanGAP B-granules have similar expression profiles to Importin β-1 B-granules: they are absent from Purkinje cells of the cerebellum neurons (white arrows, **B**), present in neurons as identified by double labeling of CA1 and cortex with RanGAP (green, white arrows) and NeuN (red, **C**), absent from neonatal cortical neurons at P1 **(D)** and primary cortical neurons labeled with RanGAP (green) and β-III-tubulin (red) at 7DIV **(E)**. Data are mean±SEM. One-way ANOVA with Bonferroni post-hoc testing did not reveal any significant differences between *C9orf72^+/-^* and *C9orf72^-/-^* mice at 2 or 6 months of age. Scale bars = 20µm. DIV = days *in vitro*, P1 = postnatal day 1.

**Figure S6**

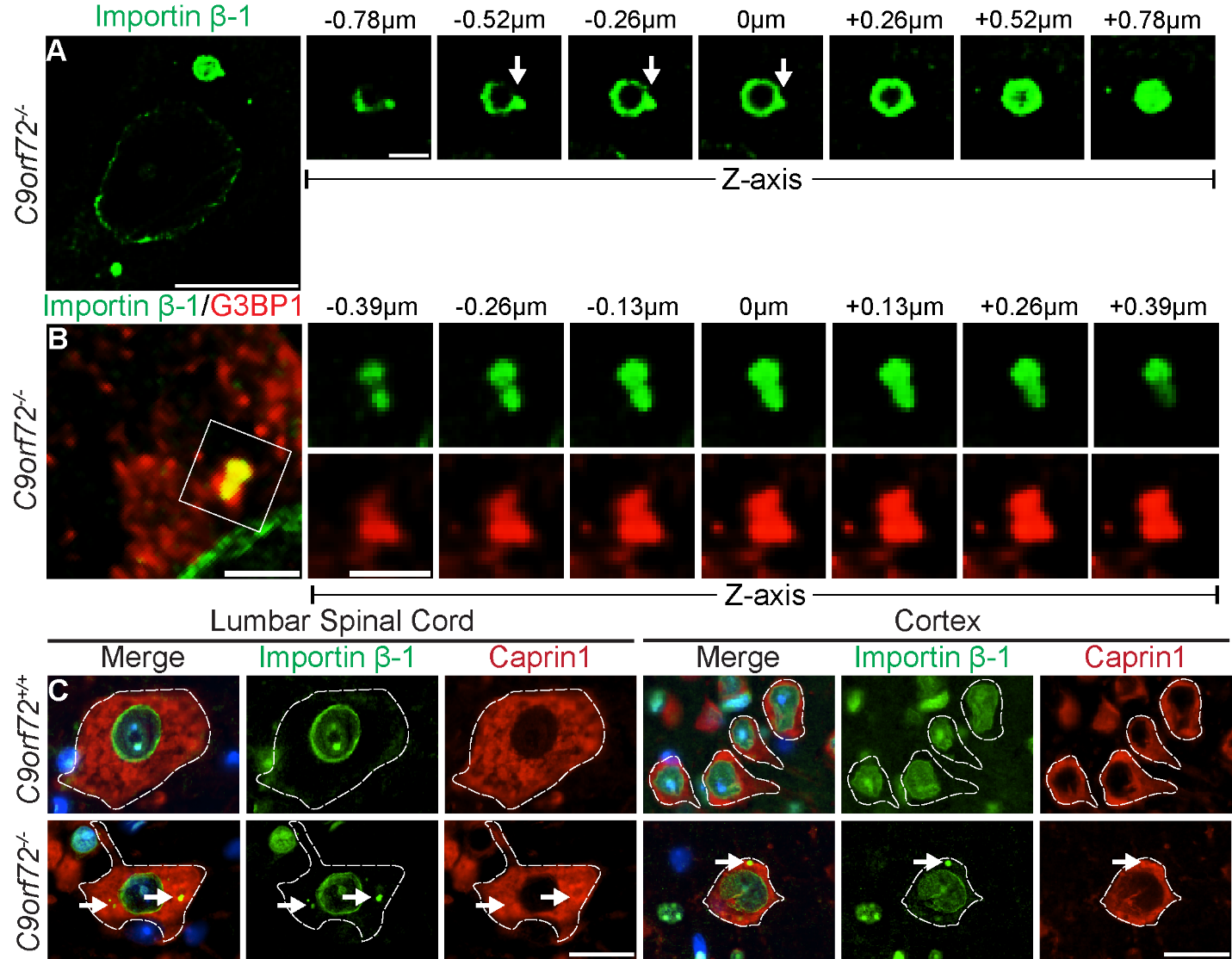

***S6: Characteristics of G3BP1 colocalization to Importin β-1 granules***

**(A)** Immunofluorescence of motor neurons labelling demonstrates that Importin β-1 is absent from the core of L-granules, which can show protuberances (white arrows) through z-slices.

**(B)** Double immunofluorescence labeling of *C9orf72^-/-^* motor neurons demonstrates that G3BP1 (red) can colocalize with Importin β-1 (green) in L-granules that appear to be merging/dividing. **(C)** Double labelling for Importin β-1 (green) and stress granule protein Caprin1 (red) does not reveal any colocalization of Caprin1 to Importin β-1 granules in lumbar spinal cord motor neurons or cortical neurons. DAPI nuclear stain (blue). Scale bars A = 10µm, 3µm panel. Scale bar B-D = 3µm, 1.5µm panels. Scale bars E = 20µm.

**Figure S7**

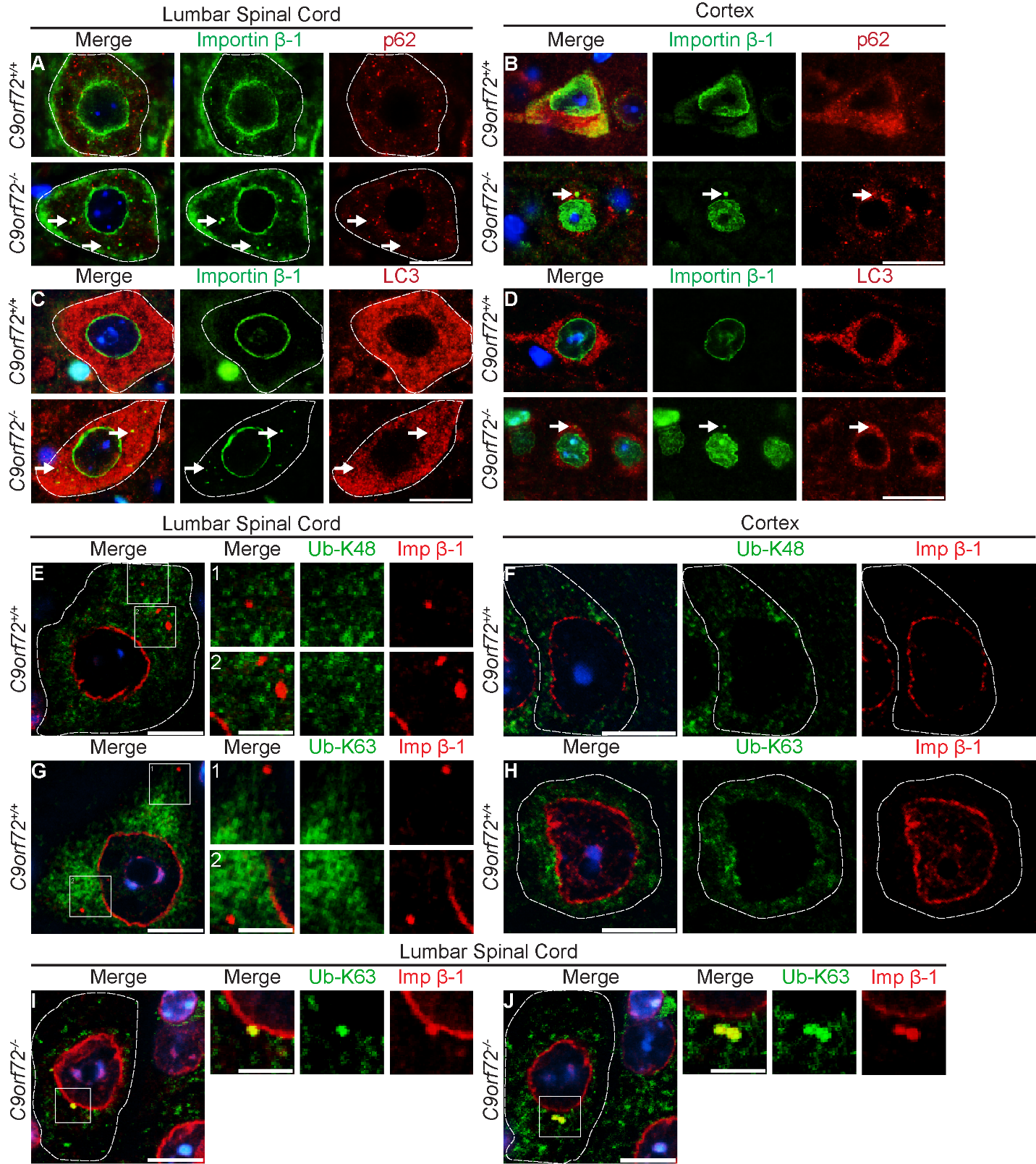

***S7: Examination of colocalizations of Importin β-1 granules with components of autophagic and ubiquitin-proteasome systems.***

**(A-D)** Double immunofluorescence labelling does not show colocalization of Importin β-1 (green) with either p62 (red, **A and B**) or LC-3 (red, **C and D**) in motor neurons or cortical neurons, at 6 months of age. **(E-H)** Double immunofluorescence for Importin β-1 (red) and ubiquitin chains linked by specific lysine residues does not show any colocalization of Importin β-1 granules with either ubiquitin linked by K48-residues (Ub-K48, green, **E and F**) or K63-residues (Ub-K63, green, **G and H**) in *C9orf72^+/+^* motor neurons or cortical neurons. **(I and J)** Double immunofluorescence labelling for ubiquitin chains linked by Ub-K63 (green) and Importin β-1 (red) in *C9orf72^-/-^* motor neurons demonstrates colocalization during granule budding **(I)** and in granules which appear to be merging or dividing **(J)**. DAPI nuclear stain (blue). Scale bars A-D= 20µm. Scale bars E-J = 10µm, 5µm in panels. Imp β-1 = Importin β-1.
